# Extreme Heterogeneity in Genomic Differentiation between Phenotypically Divergent Songbirds: A Test of Mitonuclear Co-introgression

**DOI:** 10.1101/2021.08.08.455564

**Authors:** Ellen Nikelski, Alexander S. Rubtsov, Darren Irwin

**Author notes:** Department of Ecology and Evolutionary Biology, University of Toronto, Toronto, ON, Canada. Corresponding Author: Ellen Nikelski, Department of Ecology and Evolutionary Biology, University of Toronto, Toronto, ON, Canada.

## Abstract

Comparisons of genomic variation among closely related species often show more differentiation in mitochondrial DNA (mtDNA) and sex chromosomes than in autosomes, a pattern expected due to the differing effective population sizes and evolutionary dynamics of these genomic components. Yet, introgression can cause species pairs to deviate dramatically from general differentiation trends. The yellowhammer (*Emberiza citrinella*) and the pine bunting (*E. leucocephalos*) are hybridizing avian sister species that differ greatly in appearance and moderately in nuclear DNA, but that show no mtDNA differentiation. This mitonuclear discordance is best explained by adaptive mtDNA introgression—a process that can select for co-introgression at nuclear genes with mitochondrial functions (mitonuclear genes). To better understand the extent of mitonuclear discordance and characterize nuclear differentiation patterns in this system, we investigated genome-wide differentiation between allopatric yellowhammers and pine buntings and compared it to what was seen previously in mtDNA. We found significant nuclear differentiation that was highly heterogeneous across the genome, with a particularly wide differentiation peak on the sex chromosome Z. We further tested for preferential introgression of mitonuclear genes and found statistical support for this process in yellowhammers. A role for mitonuclear coevolution in this system is supported by a stronger signal of co-introgression in genes coding for subunits of the mitoribosome and electron transport chain complexes. Altogether, our study emphasizes the extreme variation seen in differentiation across genomic components and study systems as well as highlights the ramifications of this variation in species evolution.

## Introduction

Evolution in eukaryotes is shaped by changes in multiple genomic components that differ in their modes of inheritance: mitochondrial DNA (mtDNA) is usually inherited through the matrilineal line, autosomes are inherited through both parental lines and sex chromosomes are inherited differentially depending on the sex of both parent and offspring (Avise, 2000). During speciation, populations of a single species diverge genetically, often in isolation, with the strength and pattern of genetic differentiation varying across the different genomic components (reviewed in Coyne & Orr, 2004; reviewed in Price, 2008). This variation arises as a result of differences in each component’s rate of evolution as well as the degree to which each component contributes towards reproductive isolation and is resistant to gene flow between populations in secondary contact. Most commonly, speciating taxa will show clear differentiation in mtDNA (eg. Hebert et al. 2004; Kerr et al. 2007), moderate differentiation in sex chromosomes (eg. Thornton & Long, 2002; Borge et al. 2005; Lu & Wu, 2005; Harr, 2006; Ruegg et al. 2014; Sackton et al. 2014), and comparatively modest differentiation across autosomes (Harr, 2006; Nadeau et al. 2012; Irwin et al. 2018).

In most cases, mtDNA shows strong differentiation between speciating populations (e.g. Hebert et al. 2004; Kerr et al. 2007). These patterns are partially driven by the mitochondrial genome’s uniparental inheritance and haploid nature which decrease its effective population size to ¼ that of autosomal DNA (Moore, 1995). Combined with a relatively high mutation rate (Lynch et al. 2006), this low effective population size leads to the rapid fixation of mitochondrial mutations via genetic drift and strong mtDNA differentiation between taxa. Additionally, a lack of recombination across the mitochondrial genome can further contribute to mtDNA divergence through rampant genetic hitchhiking (Hill, 2020). Here, positive selection for an adaptive mtDNA mutation causes the fixation of genetic variants across the mitochondrial genome resulting in strong differentiation between speciating taxa.

Sex chromosomes—specifically the Z (Borge et al. 2005; Ruegg et al. 2014; Sackton et al. 2014) and X chromosomes (Thorton & Long, 2002; Lu & Wu, 2005; Harr, 2006)—show moderate levels of genetic differentiation between speciating populations, but often much greater genetic differentiation than autosomes. This trend is driven by two important characteristics of Z/X chromosomes. First, on Z/X chromosomes, beneficial recessive mutations are immediately exposed to selection in the heterogametic sex allowing these advantageous variants to fix more rapidly than they would on autosomes—a phenomenon known as the “faster Z/X effect” (Meisel & Connallon, 2013; Irwin, 2018). Second, because Z/X chromosomes are inherited as either one or two copies depending on the sex, these chromosomes have a lower effective population size than autosomes (Mank et al. 2010; Irwin, 2018). This lower effective population size allows for the fixation of a greater number of neutral and slightly deleterious mutations due to less effective purifying selection and a larger role of genetic drift. Working in tandem, these two characteristics drive genetic divergence of Z/X chromosomes between speciating populations.

Across autosomes, genetic differentiation between speciating taxa is relatively modest compared to mtDNA and sex chromosomes and, interestingly, it is often characterized by “islands of differentiation”—peaks of high relative differentiation that appear within a background of low relative differentiation (Harr, 2006; Nadeau et al. 2012; Hejase et al. 2020). Explanations for these “islands” often invoke reduced gene flow between speciating taxa during secondary contact (Wu, 2001) and/or repeated bouts of selection prior to and following secondary contact (Cruickshank and Hahn, 2014; Irwin et al. 2018). In the former scenario, differentiation peaks are hypothesized to house the loci responsible for reproductive barriers between taxa making them resistant to the homogenizing influence of gene flow (Wu, 2001). In the latter scenario, differentiation islands are described as genomic areas that have experienced recurrent selection or selective sweeps (Cruickshank and Hahn, 2014; Irwin et al. 2018). These events reduce genetic diversity first in the ancestral population and then in both daughter populations producing relative peaks in genetic differentiation between taxa.

Despite the general differentiation trends discussed above, an increasing number of studies have reported differentiation patterns that greatly diverge from what is normally seen between speciating populations (Irwin et al. 2009; Yannic et al. 2010; Bryson et al. 2012). In these studies, genomic components that are normally highly divergent between taxa show remarkably low differentiation when compared to other genomic components and observable phenotypes. In particular, cases of “mitonuclear discordance”, where mtDNA shows dramatically low differentiation compared to nuclear DNA, have been identified across lineages (Toews & Brelsford, 2012). One hypothesis that may explain mitonuclear discordance is introgression of mitochondrial haplotypes from one population into another following secondary contact and hybridization. Introgression of mtDNA can occur neutrally due to various processes such as sex-biased dispersal or hybrid zone movement (Toews & Brelsford, 2012); however, the extreme degree of discordance observed in certain systems (e.g. Alves et al. 2008; Irwin et al. 2009), suggests that this process may also be driven by selection and occur adaptively by providing a fitness advantage to the receiving population.

Introgression of foreign mtDNA is thought to provide fitness advantages through two major avenues. First, because variation in mtDNA has been associated with variation in mitochondrial efficiency under different abiotic conditions (e.g. Ballard et el. 2007), mtDNA introgression may allow the receiving population to better adapt and survive within a changing or novel environment (Hulsey et al. 2016; Sloan et al. 2017; Hill, 2019). For example, mtDNA introgression has been tentatively associated with thermal adaptation in populations of rabbits (Alves et al. 2008) and cichlids (Hulsey et al. 2016). The second way that mtDNA introgression can provide a fitness advantage to a receiving population is by replacing a mitochondrial genome with a high mutational load (Sloan et al. 2017; Hill, 2019). As described earlier, the low effective population size (Moore, 1995) and high mutation rate (Lynch et al. 2006) of mtDNA can result in rapid fixation of mitochondrial mutations, including deleterious mutations. This tendency combined with genetic hitchhiking of deleterious mutations (Hill, 2020), can lead to a high mtDNA mutational load which may decrease mitochondrial efficiency. Through introgression of a foreign mitochondrial haplotype with a lower mutational load, the receiving population may be able to regain greater mitochondrial function which translates into higher fitness (Llopart et al. 2014; Hulsey et al. 2016; Sloan et al. 2017; Hill, 2019).

Adaptive mtDNA introgression presents a compelling hypothesis for the appearance of extreme “mitonuclear discordance” between speciating taxa. However, this idea becomes complicated when we consider recent work that suggests strong coevolution between the mitochondrial and nuclear genomes (Hill, 2019). In most bilaterian animals, the mitochondrial genome has been reduced to 37 genes. As a result of this low gene content, all mitochondrial processes—including oxidative phosphorylation and transcription, translation and replication of mtDNA—are reliant on more than 1000 proteins encoded by specific “mitonuclear genes” across the nuclear genome (Calvo & Mootha, 2010; Lotz et al. 2014). Tight associations between mitochondrial and mitonuclear proteins imply tight coevolution between the mitochondrial and nuclear genomes due to selection for mitochondrial efficiency: changes in one create selective pressure for compatible changes in the other (Gershoni et al. 2009; Burton & Barreto, 2012; Hill, 2019). This association also suggests that hybridization and gene flow between speciating populations will be selected against due to recombination of coevolved mitochondrial and mitonuclear genes exposing genetic incompatibilities (mitonuclear incompatibilities) within hybrid individuals. In this way, mitonuclear coevolution has the potential to contribute towards reproductive isolation between taxa and to strongly select against mtDNA introgression.

Nevertheless, if mtDNA introgression is highly adaptive such that its fitness advantages outweigh the fitness disadvantages of mitonuclear incompatibilities, this process may still be able to occur. In this scenario, researchers hypothesize that mtDNA introgression will select for co-introgression of mitonuclear genes to maintain coevolved genotypes and optimum mitochondrial function (Sloan et al. 2017; Hill, 2019). Evidence for mitonuclear co-introgression has been found in some systems (Beck et al. 2015; Morales et al. 2018; Wang et al. 2021), but the significance of this process in genome-wide nuclear differentiation and general speciation trends is still up for debate.

The yellowhammer (Passeriformes: Emberizidae: *Emberiza citrinella*) and pine bunting (*E. leucocephalos*) system (Figure 1) is one that may capture the complex interplay between genetic differentiation and mitonuclear coevolution. Thought to have diverged in isolation during the Pleistocene glaciations (Irwin et al. 2009), this Eurasian avian sister pair is highly divergent in plumage and moderately divergent in song and ecology (Panov et al. 2003; Rubtsov & Tarasov, 2017). Yet, despite their differences, yellowhammers and pine buntings hybridize extensively in a large and seemingly expanding secondary contact zone in central and western Siberia (Panov et al. 2003; 2007; Rubtsov, 2007; Irwin et al. 2009; Rubtsov & Tarasov, 2017).

**Figure 1.**
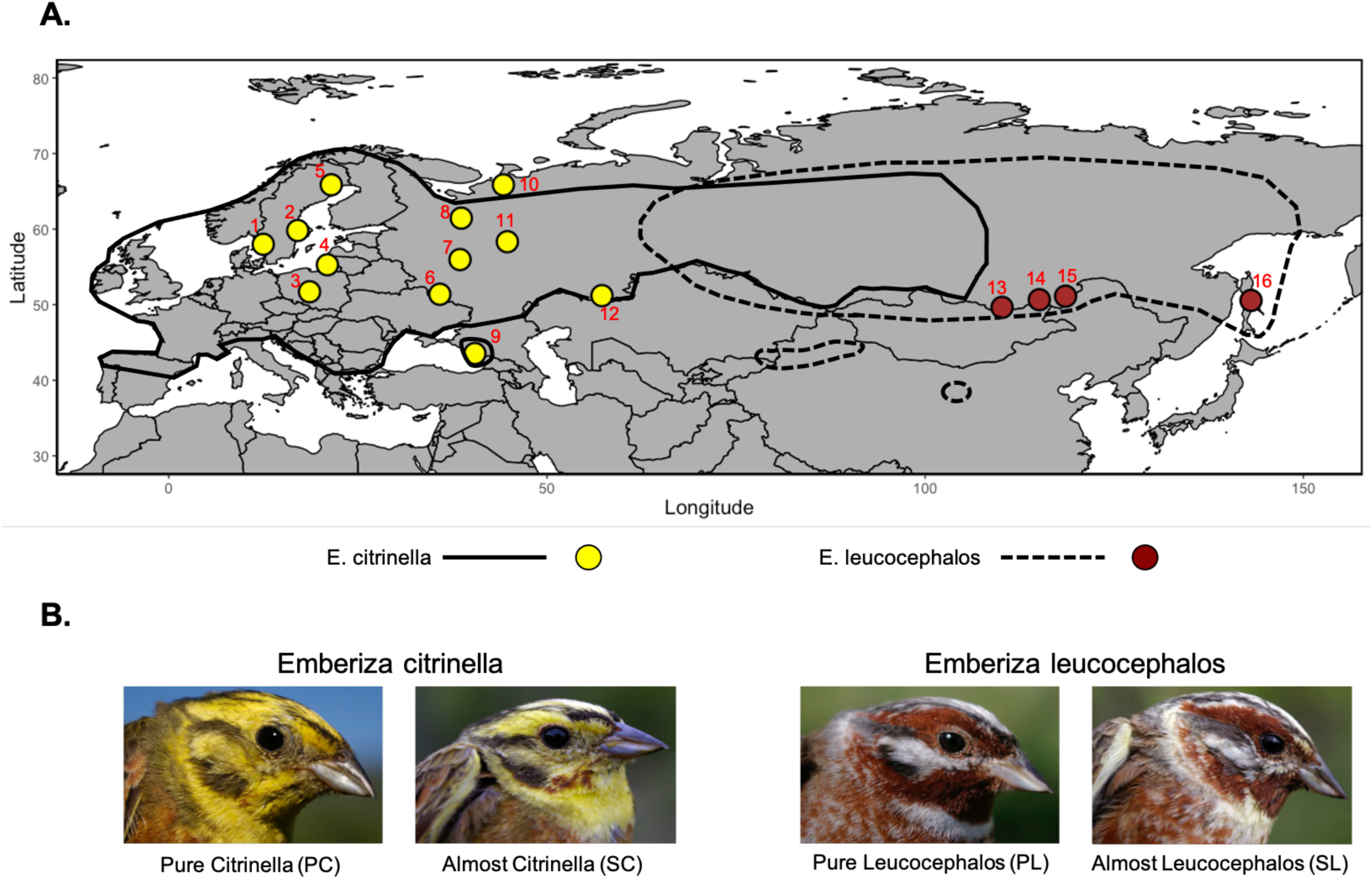
A) Map of sampling locations included in this study. Red numbers accompanying each location correspond to the sampling location numbers appearing in Table 1 which also describes sample sizes. Sampling locations may include multiple sites that appeared too close together to be shown in detail in this figure. Full details for the sites included in each sampling location can be found in Supplementary Table Sampling location points are coloured based on the taxon caught in each area: yellowhammer (*Emberiza citrinella*; yellow) and pine bunting (*Emberiza leucocephalos*; brown). The solid black line indicates the geographic range of the yellowhammer and the dashed black line indicates the geographic range of the pine bunting as described in Irwin et al. (2009). **B)** Photos of plumage variation between yellowhammers and pine buntings. Each photo represents one of four phenotypic classes: PC, SC, PL and SL. Individuals with a PC and SC phenotypic class were grouped together as *Emberiza citrinella* and individuals with a PL and SL phenotypic class were grouped together as *Emberiza leucocephalos*. All photos are credited to Dr. Alexander Rubtsov.

**Table 1.**
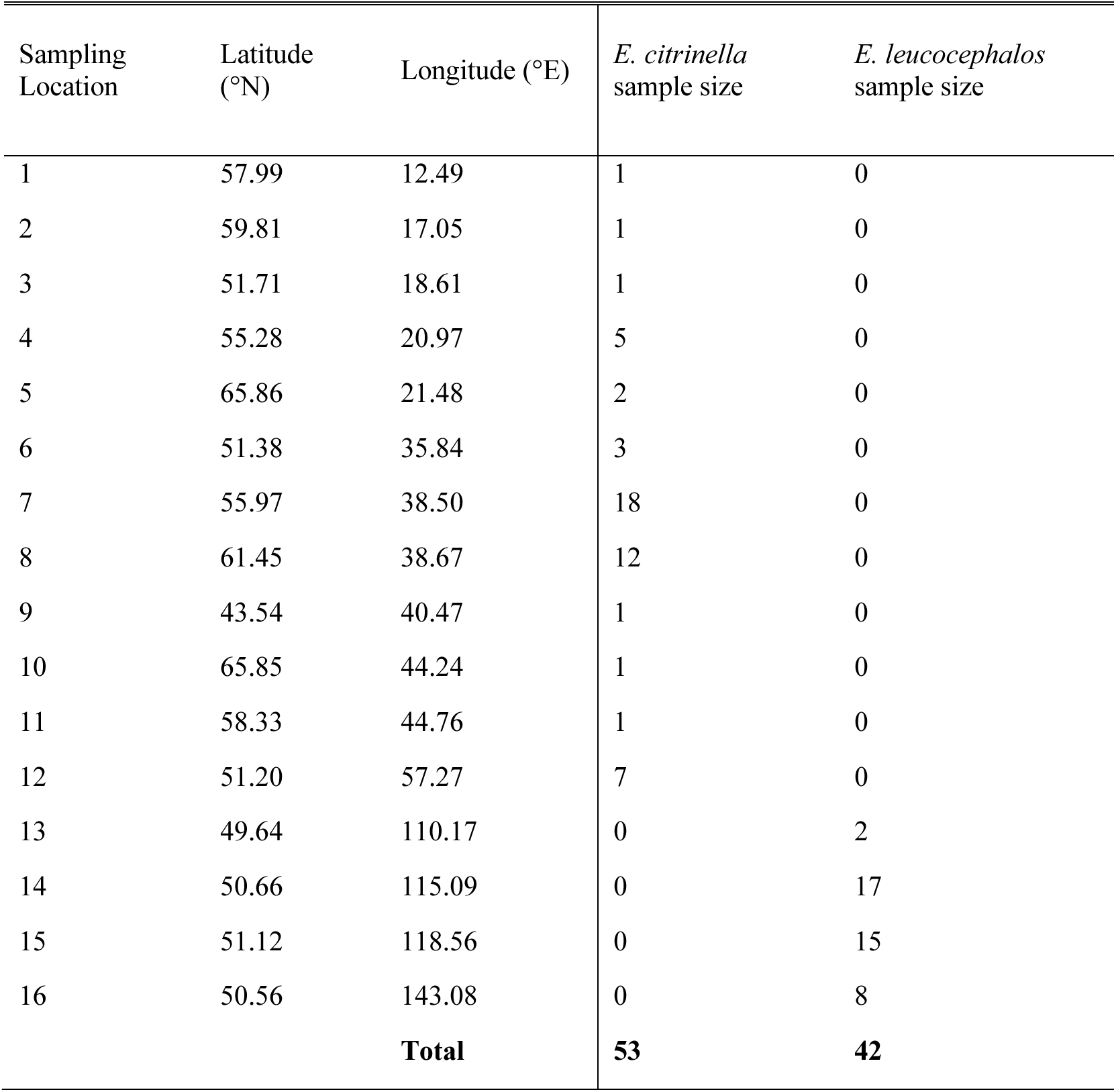
Geographic locations and sample sizes of the sites included in this study. Sampling locations may include multiple sites that appeared too close together to be shown in detail in Figure 1A. Full details for the sites included in each sampling locations can be found in Supplementary Table 1. The sampling location numbers that appear in the “Sampling Location” column correspond to those that appear in red in Figure 1A. The “Sample Size” columns describes the total number of samples collected from a particular location.

Previous genomic work has identified mitonuclear discordance between allopatric yellowhammers and pine buntings (Irwin et al. 2009) as they are nearly identical in mtDNA but show moderate differentiation in nuclear AFLP (Amplified Fragment Length Polymorphism) markers. To explain these results, Irwin et al. (2009) suggested that mtDNA introgressed adaptively from one species into the other during a previous selective sweep, and this hypothesis was supported by several statistical tests performed on the mtDNA haplotype network. Based on mitonuclear theory, such mtDNA introgression could select for similar introgression at mitonuclear genes to maintain optimum mitochondrial function (Sloan et al. 2017; Hill, 2019). The resulting lack of mitonuclear incompatibilities between taxa due to this co-introgression could facilitate their continued hybridization and hinder the build-up of reproductive barriers.

With this opposition between the observed strong nuclear differentiation and the potential for mitonuclear co-introgression, the fate of the yellowhammer and pine bunting system remains uncertain. Depending on which way the scales tip, yellowhammers and pine buntings may continue to diverge and speciate or they may continue to hybridize and eventually collapse into one interbreeding population.

Here, we present a largescale comparison of DNA sequence variation across the nuclear genomes of allopatric yellowhammers and pine buntings. With this data, we address key questions regarding genetic differentiation and mitonuclear coevolution in this system. First, what is the degree and structure of genetic differentiation between yellowhammers and pine buntings across the nuclear genome? Earlier AFLP analyses identified clear differentiation of nuclear markers between yellowhammers and pine buntings (Irwin et al. 2009), but those results were not based on actual DNA sequences and only captured a small portion of the nuclear genome. Comparing patterns of differentiation across the nuclear genome enables better understanding of the extent of mitonuclear discordance between taxa and of which genomic regions or components (e.g. sex chromosomes) show particularly high differentiation. Second, is there an over-representation of known mitonuclear genes within genomic regions that show the strongest signals of being introgressed–a pattern consistent with mitonuclear co-introgression? Evidence of mitonuclear co-introgression and the resulting lack of mitonuclear incompatibilities could explain the continued, extensive hybridization seen between yellowhammers and pine buntings and implicate this process as a force that counters divergence and the evolution of strong reproductive barriers between groups. By answering these questions, we hope to provide insight on the evolutionary trajectory of yellowhammers and pine buntings (i.e. whether it is one of continued population divergence or of population merging) and also to explore how the interplay between genetic differentiation and mitonuclear coevolution influences the speciation process more generally.

## Materials and Methods

### Sampling

We included 109 blood and tissue samples in this study: 53 phenotypic yellowhammers, 42 phenotypic pine buntings, and 14 other members of Emberizidae (one *Emberiza aureola* [yellow-breasted bunting], one *Emberiza calandra* [corn bunting], one *Emberiza cioides* [meadow bunting], one *Emberiza hortulana* [ortolan bunting], four *Emberiza stewarti* [white-capped bunting], and six *Emberiza cirlus* [cirl bunting]) to put variation between yellowhammers and pine buntings into a deeper phylogenetic context (Figure 1A; Table 1; Supplementary Table 1). A total of 91 of our samples were included in the AFLP analysis of Irwin et al. (2009) while 18 samples were examined for the first time as part of the present research.

When possible, body measurements and photographs were taken of live birds or museum skins. Yellowhammer and pine bunting males were scored phenotypically and sorted into phenotypic classes based on the protocols presented in Panov et al. (2003) and Rubtsov & Tarasov (2017). Briefly, each male received a score from 0-7 for three plumage traits, where 0 represents a phenotypically pure yellowhammer and 7 represents a phenotypically pure pine bunting. These three traits were (1) the amount of chestnut plumage (vs. yellow or white) at the brow, (2) the amount of chestnut plumage (vs. yellow or white) at the throat, and (3) background plumage colour, meaning the strength of yellow—ranging from bright yellow to pure white—in head and body plumage that did not show brown or black streaking. Phenotypic classes included: pure *citrinella* (PC), almost *citrinella* (SC, for “semi-*citrinella*”), *citrinella* hybrid (CH), yellow hybrid (YH), white hybrid (WH), *leucocephalos* hybrid (LH), almost *leucocephalos* (SL, for “semi-*leucocephalos*”) and pure *leucocephalos* (PL) (Rubtsov & Tarasov, 2017). Any SC and SL individuals that appeared in the allopatric zones were grouped together with PC and PL individuals respectively and treated as phenotypic yellowhammers and phenotypic pine buntings in subsequent analyses (Figure 1B).

### DNA extraction and genotyping-by-sequencing

DNA was extracted from samples using a standard phenol-chloroform method. We then divided the DNA samples into four genotyping-by-sequencing (GBS) libraries (Elshire et al. 2011). The 109 samples included in this study were sequenced together with 226 yellowhammer, pine bunting and hybrid DNA samples collected near and within the sympatric zone as part of a larger project (Nikelski et al. in prep). The libraries were prepared as per the protocol described by Alcaide et al. (2014) with modifications specified by Geraldes et al. (2019) except that we maintained a 300-400 bp fragment size during size selection. Paired-end sequencing was completed by Genome Québec using an Illumina HiSeq 4000 system, producing more than 1.2 billion reads, each 150 bp in length, across the four GBS libraries.

### Genotyping-by-sequencing data filtering

We processed the reads following Irwin et al. (2016; 2018), as summarized here. Reads were demultiplexed using a custom perl script designed by Baute et al. (2016). Next, reads were trimmed for quality using Trimmomatic version 0.36 (Bolger et al. 2014) with the parameters: TRAILING:3, SLIDINGWINDOW:4:10, MINLEN:30. Trimmed reads were aligned to the zebra finch reference genome (*Taeniopygia guttata* version 3.2.4; Warren et al. 2010) using the program BWA-MEM (Li & Durbin, 2009) and a BAM file of this information was created for each individual using the programs Picard (http://broadinstitute.github.io/picard/) and SAMtools (Li et al. 2009). BAM files were converted into GVCF files using the HaplotypeCaller command as part of GATK version 3.8 (McKenna et al. 2010). We then combined information from the individuals in two ways to create 1) a genome-wide “variant site” VCF file containing only variant site information, and 2) a series of chromosome-specific “info site” VCF files containing information on both variant and invariant sites with sufficient coverage.

To create the genome-wide “variant site” VCF file, we used the GenotypeGVCFs command in GATK to identify and isolate single nucleotide polymorphisms (SNPs) among individuals. This command also converted the variant site information into a single VCF file encompassing the entire nuclear genome. Using a combination of VCFtools (Danecek et al. 2011) and GATK, we filtered the VCF file to remove indels and non-biallelic SNPs. We also discarded loci with QD < 2.0, MQ < 40.0, FS > 60.0, SOR > 3.0, or ReadPosRankSum < -8.0. Finally, loci with more than 60% missing genotypes were removed. The average coverage of variable sites in the resulting VCF file was 16.59.

To convert GVCF files into “info site” VCF files, we similarly employed the GenotypeGVCFs command in GATK with the addition of the -allSites and -L flags to retain invariant sites and split the information into chromosome-specific files. The resulting VCF files were filtered using VCFtools and GATK to remove indels, sites with more than two alleles, sites with more than 60% missing data, sites with MQ values lower than 20 and sites with heterozygosities greater than 60% (to avoid potential paralogs).

### Variant site analyses

The genome-wide “variant site” VCF file was analyzed using modified versions of the R scripts described in Irwin et al. (2018), and all of our analyses used R version 3.6.2 (R Core Team, 2014). A total of 374,780 SNPs were identified among allopatric yellowhammers and pine buntings. For each of these SNPs, we calculated sample size, allele frequency, and Weir and Cockerham’s *F*_ST_ (Weir & Cockerham, 1984). Genetic differentiation between yellowhammers and pine buntings was then visualized using a principal components analysis (PCA) generated with the pca command and the svdImpute method to account for any missing genomic data using the pcaMethods package (Stacklies et al. 2007). A Manhattan plot of *F*_ST_ values for 349,807 SNPs with known genomic locations was created using the package qqman (Turner, 2018).

### Differentiation across the genome

To thoroughly investigate nuclear differentiation between allopatric yellowhammers and pine buntings, we performed further analysis on both variant and invariant loci within “info site” VCF files using R scripts described in Irwin et al. (2018).

We calculated Weir and Cockerham’s *F*_ST_, between-group nucleotide differentiation (*π*_B_) and within-group nucleotide variation (*π_w_*) within nonoverlapping windows of available sequence data across each chromosome. The first window was positioned at the “start” of each chromosome as described in the zebra finch reference genome (Warren et al. 2010). We used a window size of 2,000 bp of sequenced data rather than 10,000 bp (as in Irwin et al. 2018), to visualize narrow peaks in relative and absolute differentiation within our dataset. We hereafter refer to these windows as “genomic windows.”

We developed a new R script to calculate a Tajima’s D value (Tajima, 1989) for each of the genomic windows. Values of Tajima’s D were used to identify areas of the genome where patterns of variation in yellowhammer and pine bunting populations deviated from a neutral model. Significantly negative Tajima’s D implies that the ratio of common vs. rare alleles is lower than expected under neutrality, likely because of a selective sweep or population expansion following a bottleneck. Significantly positive Tajima’s D suggests that the ratio of common vs. rare alleles is higher than expected under neutrality, potentially stemming from balancing selection or a rapid population contraction.

### Phylogenetic comparison with other Emberizidae species

We employed whole-genome averages of *π*_B_ between allopatric yellowhammers and pine buntings as well as among these focal species and six other Emberizidae species (*Emberiza aureola, Emberiza calandra, Emberiza cioides, Emberiza cirlus, Emberiza hortulana and Emberiza stewarti*) to estimate a phylogeny. A list of average *π*_B_ values for each species pair was converted into a distance matrix and used to create an unrooted neighbour-joining tree. This tree was constructed using the ape package (Paradis & Schliep, 2019) and the BioNJ algorithm (Gascuel, 1997) with *Emberiza aureola* set as the outgroup (Alström et al. 2008). The phylogeny was created to provide further support for the sister relationship between yellowhammers and pine buntings that had previously been hypothesized using mitochondrial markers (Alström et al. 2008; Irwin et al. 2009) but that was questioned in some studies (Rubtsov & Opeav, 2012). In creating this phylogeny we were also able to better investigate the degree of mitonuclear discordance within the system using a greater amount of nuclear data.

### Signals of mitonuclear co-introgression

To test for preferential mitonuclear gene introgression (compared to other genomic regions) between allopatric yellowhammers and pine buntings, we asked whether there was an association between a list of known mitonuclear genes and a list of genomic windows that have statistical characteristics most consistent with introgression. We call the latter “putative introgression windows” (PIWs), and we explain their identification in detail below.

The mitonuclear genes that we analyzed for signals of introgression were all protein coding with products that interacted directly with mtDNA or an immediate product of the mitochondrial genome (i.e. protein or RNA). For these nuclear-encoded genes, any change in mtDNA including those caused by introgression would likely cause selection for co-introgression of compatible alleles (Gershoni et al. 2009; Burton & Barreto, 2012; Hill, 2019). Mitonuclear genes that met these criteria included those that encode protein subunits of ATP synthase or the first, third and fourth complex of the electron transport chain (ETC), assembly and ancillary proteins involved in the formation of the ETC, and proteins that are part of the transcription, translation or DNA replication machinery within the mitochondria (Diodato et al. 2013; Greber and Ban 2016; Hill, 2019). After removing any genes that were not annotated in the zebra finch reference genome or that lacked a specific location on the reference genome, a total of 162 mitonuclear genes remained for analysis (Supplementary Table 2).

Given the ongoing hybridization observed between yellowhammers and pine buntings and the evidence for mtDNA introgression and replacement, it is reasonable to assume that regions of the nuclear genome have also moved between the species. For each taxon, we identified a subset of genomic windows that showed statistical properties most consistent with substantial introgression, although we cannot be certain of introgression for any particular PIW. We identified PIWs as those that possessed a low *π_B_* average and a low Tajima’s D average. Low *π_B_* indicates high similarity between the nucleotide sequences of the two groups as would be expected if alleles had introgressed from one taxon into the other. Low Tajima’s D suggests a past selective sweep within a population which would also be expected if an adaptive allele had introgressed from a separate taxon and swept throughout the receiving population. For this analysis, our quantitative criteria for a PIW were a Tajima’s D value within the lowest 5% of the available windowed values and a *π_B_* value within the lowest 30% of the available windowed values. The Tajima’s D threshold was kept relatively low to capture windows with particularly strong signals of selection that could be associated with adaptive introgression while the *π*_B_ threshold was left higher to identify an appreciable number of PIWs for analysis.

Using a custom R script, each mitonuclear gene was assigned to the genomic window that minimized the absolute difference between the location of the mitonuclear gene centre and the location of the window centre. We then determined the number of mitonuclear genes that occurred within the PIWs of each taxon.

We conducted a Fisher’s Exact test for both yellowhammers and pine buntings to determine whether the proportion of mitonuclear genes within PIWs was significantly different from what would be expected based on the total proportion of protein coding genes appearing within these windows. To complete this analysis, a list of the 14,008 protein-coding genes annotated in the zebra finch reference genome (Warren et al. 2010) was compiled. We removed our 162 mitonuclear genes from this list and assigned the remaining 13,846 non-mitonuclear genes to genomic windows using the methodology described above. The proportion of mitonuclear and non-mitonuclear genes appearing in PIWs were then compared separately in yellowhammers and pine buntings.

## Results

When comparing allopatric yellowhammers and pine buntings, we identified 374,780 variable SNPs within our “variant site” VCF file and 13,703,455 invariant and 699,122 variant sites across thirty autosomes and the Z chromosome within our “info site” VCF files following filtering. In the latter “info site” files, we designated a total of 7,187 genomic windows (of 2000 sequenced bp each) across the genome, with each window covering an average distance of about 139 kilobases.

### Phylogenetic comparison with other Emberizidae species

An unrooted neighbour-joining tree of average *π_B_* values between yellowhammers, pine buntings and six other Emberizidae species (Figure 2) depicted similar species relationships as were identified previously using mitochondrial markers (Alström et al. 2008; Irwin et al. 2009). Relative branch lengths were also similar, with the major exception being the branch length between yellowhammers and pine buntings which was much longer in our analysis using nuclear DNA. To put this into context, we determined the relative genetic distance between yellowhammers and pine buntings versus the genetic distance between *E. stewarti* and either member of the yellowhammer/pine bunting clade for our nuclear phylogeny and for the previously calculated mitochondrial phylogeny (Irwin et al. 2009). The nuclear ratio was 11.4 times greater than the mitochondrial ratio which corroborates the presence of strong mitonuclear discordance between yellowhammers and pine buntings. These results also support the hypothesis of an extended period of divergence between yellowhammers and pine buntings followed by adaptive mtDNA introgression.

**Figure 2.**
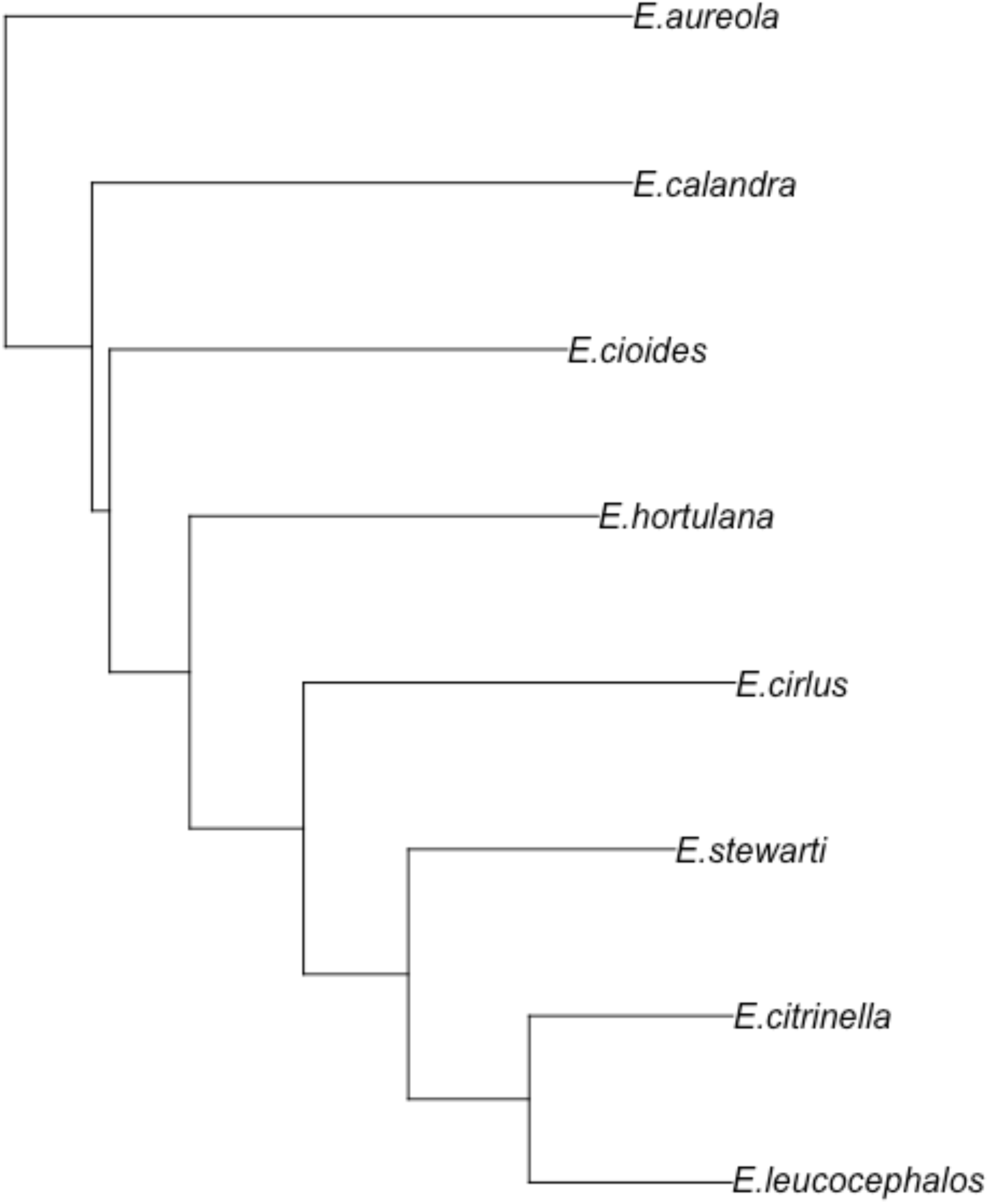
Unrooted neighbour-joining tree of Emberizidae species constructed based on average absolute between-population nucleotide diversity (*π_B_*). Sample sizes for each species are as follows: *E. aureola* = 1, *E. calandra* = 1, *E. cioides* = 1, *E. hortulana* = 1, *E. cirlus =* 6, *E. stewarti* = 4, *E. citrinella* = 53 and *E. leucocephalos* = 42.

### Overall genetic differentiation

Based on 374,780 SNPs, the genome-wide *F*_ST_ estimate was 0.0232 between allopatric yellowhammers and pine buntings. Despite this low average, a PCA based on the same SNP genotypes separated yellowhammers and pine buntings into tight genetic clusters (Figure 3). Two pine buntings were outliers along PC1, while the remaining yellowhammers and pine buntings separated into distinct groups mainly along PC2. Further investigation into these outliers revealed that they were males from the same location, but a kinship analysis completed as part of a separate study did not find close kinship between the two pine buntings that could explain their position (Nikelski et al. in prep). We also examined the PC1 loadings and found that the signal for PC1 position was broadly distributed across the nuclear genome, rather than being concentrated in a few highly influential regions (Supplementary Figure 1). Finally, we temporarily removed one of the outliers and re-ran the PCA. This caused the other outlier to fall into the pine bunting cluster, but revealed a further yellowhammer outlier (Supplementary Figure 2). Removal of this yellowhammer outlier in addition to one member of the pine bunting outlier pair in turn revealed another yellowhammer outlier (Supplementary Figure 3). It is unclear what is responsible for these outliers, but the distinct yellowhammer and pine bunting genetic clusters remained intact in all the PCAs considered.

**Figure 3.**
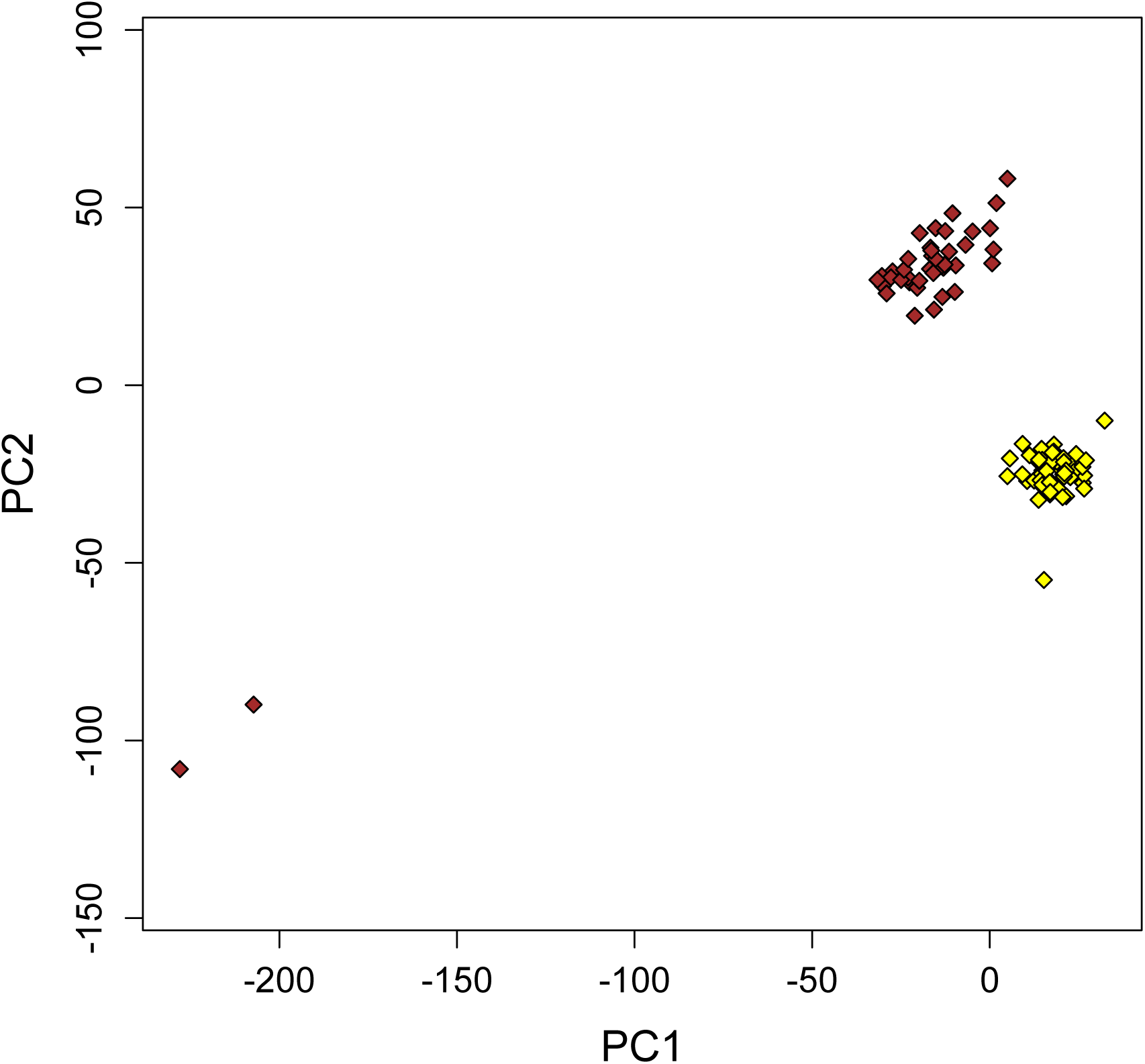
PCA of genetic variation between allopatric yellowhammers (yellow; n = 53) and allopatric pine buntings (brown; n = 42), based on 374,780 genome-wide SNPs. PC1 and PC2 explain 3.6% and 2.9%, respectively, of the variation among individuals.

### Differentiation across the genome

Relative differentiation between allopatric yellowhammers and pine buntings was highly heterogeneous across the nuclear genome with peaks in *F*_ST_ seen on most of the larger chromosomes (Figures 4, 5, 6A-B; Supplementary Figure 4). Chromosome Z in particular showed a large peak in *F*_ST_ with several SNPs possessing values close to one. In fact, *F*_ST_ for the Z chromosome was 0.1246—more than five times larger than the genome-wide *F*_ST_.

**Figure 4.**
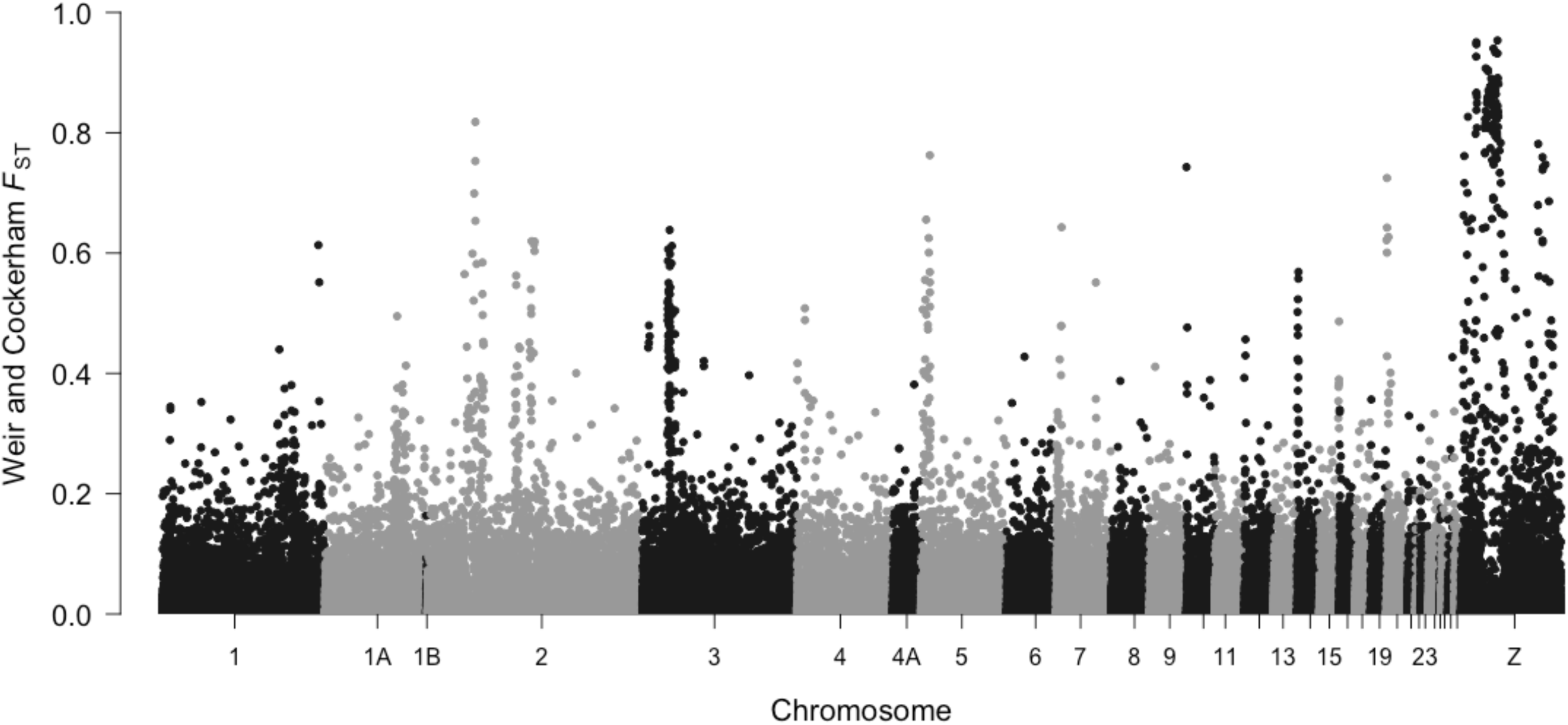
Relative differentiation (*F*_ST_) of 349,807 genome-wide SNPs identified between allopatric yellowhammers (n = 53) and allopatric pine buntings (n = 42), with chromosomes represented with alternating black and grey. Narrow regions of elevated differentiation can be seen on many autosomes, and there are broad regions of high differentiation on the Z chromosome.

**Figure 5.**
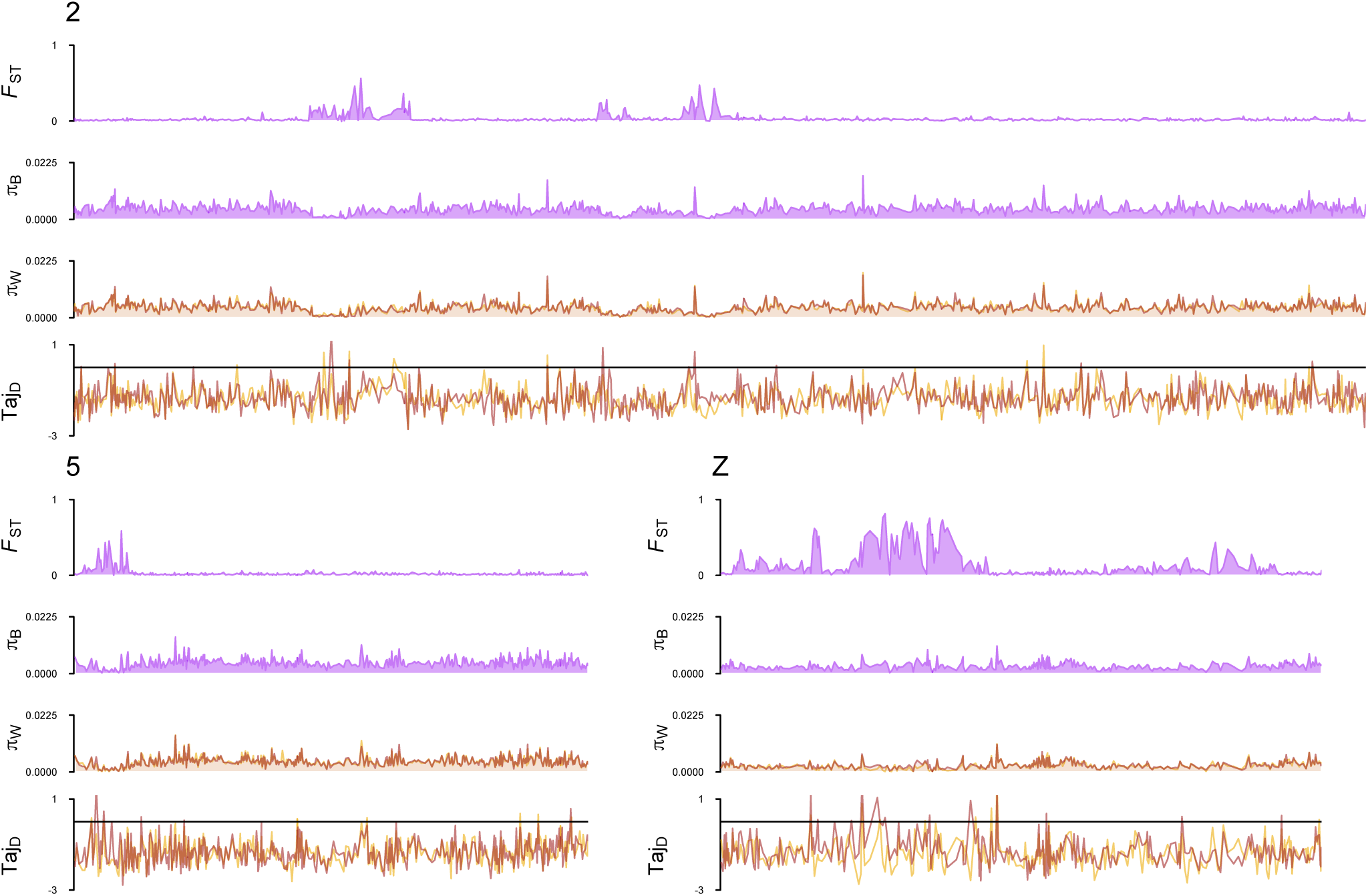
Patterns of genetic variation comparing allopatric yellowhammers (n = 53) and allopatric pine buntings (n = 42) across three chromosomes (2, 5 and Z). Relative nucleotide differentiation (*F*_ST_), absolute between-population nucleotide differentiation (*π_B_*), absolute within-population nucleotide variation (*π_W_*) and Tajima’s D (Taj_D_) are shown as 2000 bp windowed averages across each chromosome. *F*_ST_ and *π_B_* are shown as purple lines to indicate that values were calculated as a comparison between allopatric yellowhammers and pine buntings. *π_W_* and Taj_D_ are shown as two separate lines (yellow = yellowhammers, brown = pine buntings) to indicate that values were calculated separately for each population.

**Figure 6.**
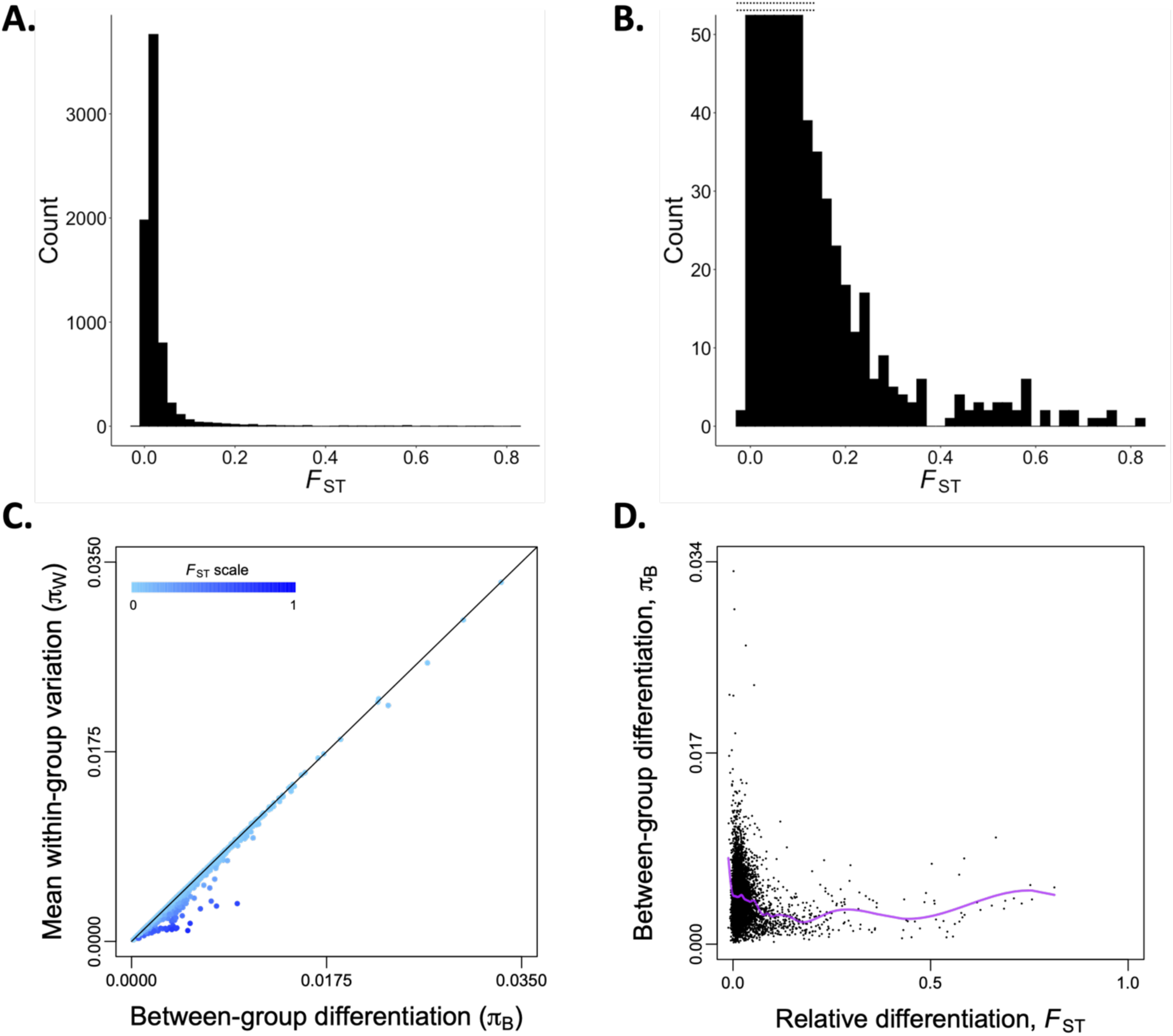
A) A histogram of average relative differentiation (*F*_ST_) values calculated for 2000 bp windows across the nuclear genome when comparing allopatric yellowhammers (n=53) with allopatric pine buntings (n=42). **B)** A truncated version of Figure 6A that depicts the high *F*_ST_ tail of the histogram. The limits of the y-axis were set as (0,50) with the double dotted line indicating bars of the histogram that have been truncated. **C)** Mean absolute within-group nucleotide variation (*π_W_*) of allopatric yellowhammers and allopatric pine buntings plotted against absolute between-group nucleotide differentiation (*π_B_*). Each dot represents the average value taken from a 2000 bp window of sequenced data across the nuclear genome. The black line indicates where mean within-group nucleotide variation equals between-group nucleotide differentiation. Increasing values of relative differentiation (*F*_ST_) calculated for each window are shown in darker shades of blue. **D)** Association between relative differentiation and absolute between-group nucleotide differenation of allopatric yellowhammers and allopatric pine buntings. Each black dot represents average values calculated from a 2000 bp window of sequenced data. A cubic spline fit between the variables is shown as a purple line.

Patterns of between-group nucleotide differentiation (*π_B_*) and within-group nucleotide variation (*π_W_*) were also heterogenous across the genome and comparable to each other in magnitude: genome-wide *π_B_* = 0.0041; genome-wide *π_w_* for both taxa= 0.0040 (Figure 5; Supplementary Figure 4). Because between-group and within-group nucleotide differentiation are so intimately related in their evolution and calculation, it is expected that windowed averages of these two statistics will show a highly positive relationship. In fact, most windowed *π_B_* and *π_w_* averages fell near a 1:1 association line (Figure 6C) which is equivalent to no or little differentiation. However, some genomic windows showed much reduced *π_w_* compared to *π_B_*; these were the windows with high *F_ST_*. Additionally, we detected a weak but highly significant negative correlation between the windowed averages of *F*_ST_ and *π_B_* (Spearman’s Rank Correlation: -0.1196, p < 2.2 × 10^-16^; Figure 6D) as is hypothesized if peaks in relative differentiation are products of repeated selective events (Cruickshank & Hahn, 2014; Irwin et al. 2018).

Finally, we found that Tajima’s D varied across the genome but was mostly negative (Figure 5; Supplementary Figure 4), consistent with a history of population growth and/or selective sweeps. The average genome-wide Tajima’s D was similar between populations: -1.377 for yellowhammers and -1.335 for pine buntings.

### Signals of mitonuclear co-introgression

Of the 7187 genomic windows identified across the nuclear genome, we classified 244 (3.4%) as PIWs within yellowhammers and 222 (3.1%) as PIWs within pine buntings. Average values of *π_B_* and Tajima’s D in yellowhammer PIWs were 0.0016 and -2.3751 respectively, and 0.0019 and -2.3369 in pine bunting PIWs respectively. In non-PIWs, the average values of *π_B_* and Tajima’s D were 0.0042 and -1.3416 in yellowhammers and 0.0042 and -1.3031 in pine buntings. Of the PIWs identified in yellowhammer and pine bunting populations, 71 were shared between the taxa. It should be noted that sharing of some PIWs is expected given that the contribution of *π_B_* to window selection was identical for both taxa (in contrast, Tajima’s D was calculated separately for yellowhammers and pine buntings).

Our examination of the gene content within yellowhammer PIWs revealed that they contained a higher percentage (7.4%) of mitonuclear genes (12 of the 162 genes considered) than of non-mitonuclear genes (4.1%; 574 of the 13,846 genes considered). This difference was statistically significant (Fisher’s Exact test: p = 0.04714), providing evidence for mitonuclear genes preferentially appearing within yellowhammer PIWs. Pine bunting PIWs contained 4.3% of the mitonuclear genes (7 of the 162 genes considered) and 3.3% of the non-mitonuclear genes (455 of the 13,846 genes considered), a difference that was not statistically significant (Fisher’s Exact test: p = 0.3806).

The twelve mitonuclear genes that appeared within yellowhammer PIWs were: APOPT1, COX5A, COX17, LARS2, MRPL1, MRPL27, MRPL32, MRPS7, MRPS25, NDUFC1, SSBP1 and UQCR11 (Table 2). Five of these genes encode protein subunits of the mitoribosome, three encode structural subunits of the ETC, two encode assembly factors of the ETC, one encodes a mitochondrial aminoacyl-tRNA synthetase and one encodes a single-stranded DNA-binding protein involved in mtDNA replication. Two genes each appear on chromosomes 2, 4 and 18 while the rest appear on separate chromosomes. Interestingly, three of the five putatively introgressed genes associated with the ETC are specifically associated with complex IV.

**Table 2.** Identities, chromosomal locations, windowed Tajima’s D values and functions of mitonuclear genes that appeared within 244 yellowhammer PIWs or within 222 pine bunting PIWs. In the “Mitonuclear Gene Function” column, ETC stands for “Electron Transport Chain”. The “*” indicates a mitonuclear gene that appeared in both yellowhammer and pine bunting PIWs. The “^†^” indicates two genes that appeared within the same PIW.

| Mitonuclear Gene | Chromosome where mitonuclear gene is found | Windowed Tajima’s D Value | Mitonuclear Gene Function |
| --- | --- | --- | --- |
| *Yellowhammer (E. citrinella)* |  |  |  |
| APOPT1 | 5 | -2.207 | Assembly factor/ancillary protein for ETC complex IV |
| COX5A* | 10 | -2.420 | Structural subunit of ETC complex IV |
| COX17 | 1 | -2.509 | Assembly factor/ancillary protein for ETC complex IV |
| LARS2 | 2 | -2.238 | Mitochondrial aminoacyl-tRNA synthetase |
| MRPL1 | 4 | -2.207 | Mitochondrial large ribosomal subunit protein |
| MRPL27 | 18 | -2.214 | Mitochondrial large ribosomal subunit protein |
| MRPL32 | 2 | -2.306 | Mitochondrial large ribosomal subunit protein |
| MRPS7* | 18 | -2.628 | Mitochondrial small ribosomal subunit protein |
| MRPS25 | 12 | -2.323 | Mitochondrial small ribosomal subunit protein |
| NDUFC1 | 4 | -2.399 | Structural subunit of ETC complex I |
| SSBP1 | 1A | -2.362 | Single stranded DNA-binding protein |
| UQCR11 | 28 | -2.499 | Structural subunit of ETC complex III |
| *Pine bunting (E. leucocephalos)* |  |  |  |
| ATP5H<sup>†</sup> | 18 | -2.601 | Structural subunit of ETC complex V |
| COX5A* | 10 | -2.545 | Structural subunit of ETC complex IV |
| MRPL2 | 3 | -2.247 | Mitochondrial large ribosomal subunit protein |
| MRPL58<sup>†</sup> | 18 | -2.601 | Mitochondrial large ribosomal subunit protein |
| MRPS7* | 18 | -2.252 | Mitochondrial small ribosomal subunit protein |
| MRPS14 | 8 | -2.173 | Mitochondrial small ribosomal subunit protein |
| NDUFB4 | 1 | -2.304 | Structural subunit of ETC complex I |

The seven mitonuclear genes that appeared within pine bunting PIWs were: ATP5H, COX5A, MRPL2, MRPL58, MRPS7, MRPS14 and NDUFB4 (Table 2). Four of these genes encode protein subunits of the mitoribosome and three encode structural subunits of the ETC. Three genes appear on chromosome 18—with two genes sharing the same genomic window— while the rest of the genes appear on separate chromosomes. The COX5A and MRPS7 genes was found in both yellowhammer and pine bunting PIWs.

## Discussion

Yellowhammers and pine buntings show negligible mtDNA differentiation (Irwin et al. 2009) but are well differentiated phenotypically (Panov et al. 2003, Rubtsov & Tarasov, 2017) and moderately differentiated in AFLP nuclear markers (Irwin et al. 2009). In the wake of this mitonuclear discordance, Irwin et al. (2009) proposed that mtDNA adaptively introgressed between taxa following a period of allopatric isolation; however, due to the limited information provided by AFLP analyses, the extent of mitonuclear discordance and of nuclear differentiation between yellowhammers and pine buntings remained unknown lending some uncertainly to this hypothesis. In the present study, analysis of genetic variation across the nuclear genome identified heterogeneous nuclear differentiation between allopatric populations with strong differentiation peaks that separated taxa into distinct genetic clusters. This result supports yellowhammers and pine buntings experiencing a period of separate evolution followed by the hybridization within their current contact zone in western and central Siberia. Our phylogenetic analysis showing a longer branch length between yellowhammers and pine buntings based on nuclear markers—when compared to a phylogeny based on mtDNA—also corroborates a hypothesis of recent mtDNA introgression and mitochondrial haplotype replacement in this system likely driven by selection (Irwin et al. 2009). Additionally, our finding that mitonuclear genes are over-represented in putative introgression windows in yellowhammers is consistent with mitonuclear co-introgression from pine buntings into yellowhammers.

Though genetically distinct, the genome-wide *F*_ST_ between allopatric yellowhammers and pine buntings (0.0232) was comparable to or sometimes lower than the averages seen between avian subspecies (e.g., subspecies of barn swallow: 0.017-0.026 [Scordato et al. 2017]; myrtle warbler and Audubon’s warbler: 0.077-0.106 [Irwin et al. 2018]; yellow- and red-shafted northern flickers: 0.098 [Manthey et al. 2016]). This low genome-wide *F*_ST_ contrasts with the moderate *F*_ST_ averages reported from an analysis of AFLP markers performed on the same populations: 0.078 based on allele frequencies and 0.140 based on band frequencies (Irwin et al. 2009). However, the present study also revealed that relative differentiation was highly heterogeneous across the nuclear genome with *F*_ST_ peaks on various chromosomes. It is possible that the previous AFLP analysis captured a disproportionate number of loci within these differentiation peaks, thereby inflating *F*_ST_ estimates. This comparison highlights the caution that should be taken when interpreting genome-wide averages as highly variable genetic differentiation landscapes can cause large variability in *F*_ST_ estimates when they are based on a limited and non-random sample of loci.

The *F*_ST_ peaks seen between yellowhammers and pine buntings on larger autosomes and most significantly on the Z chromosome are consistent with the “islands of differentiation” often noted in comparisons of closely related taxa (Harr, 2006; Nadeau et al. 2012; Irwin et al. 2018). In contrast to these islands, large regions of close similarity in *π_B_* and *π_w_* suggests high gene flow between taxa across much of the nuclear genome. This scenario is consistent with the observed extensive hybridization between these taxa (Panov et al. 2003; 2007; Rubtsov, 2007; Rubtsov & Tarasov, 2017). Nevertheless, the high *F*_ST_ islands, those with much reduced *π_w_* compared to *π_B_*, can be explained by divergent selection causing low gene flow in these regions.

It is unlikely that this pattern is the result of genetic drift over an extended period of geographic separation, as this would result in most genomic regions deviating slightly from *π_B_* = *π_w_* congruence rather than the observed pattern of extreme heterogeneity. Instead, this trend suggests that selection acted in a way that lowered *π_w_* relative to *π_B_* within “islands of differentiation”. Considering that high *F*_ST_ regions were associated with relatively low values of *π_B_*, we propose that differentiation islands in this system are most consistent with a model invoking repeated bouts of selection that lower nucleotide diversity (Cruickshank & Hahn, 2014; Irwin et al. 2018). A sweep-before-differentiation model (Irwin et al. 2018), where *F*_ST_ peaks are produced by adaptive selective sweeps between populations followed by adaptive selection at the same regions in local populations, is particularly in line with the extensive hybridization presently observed between yellowhammers and pine buntings.

Of the “islands of differentiation” identified between taxa, the tallest and widest was found on the Z chromosome. Greater differentiation on the Z chromosome compared to autosomes is a common observation when comparing closely related species (Borge et al. 2005; Ruegg et al. 2014; Sackton et al. 2014) and is consistent with stronger positive selection (the “faster Z/X effect") and with less efficient purifying selection on this chromosome (Mank et al. 2010; reviewed in Meisel & Connallon, 2013; reviewed in Irwin, 2018). However, the large region of the Z chromosome that have *F*_ST_ values near zero suggests that additional factors are involved in producing this island of differentiation.

One possible explanation for the large differentiation island on chromosome Z could be that it corresponds with an area of low recombination—a region of connected loci that tend to be inherited together, leading to linked selection of nearby loci. Strong divergent selection acting on one SNP within this region would act similarly on all the loci that are linked to it such that a wide, highly divergent genomic block would become fixed and appear as an “island” between taxa (reviewed in Cutter & Payseur, 2013). Areas of low recombination and linkage are often associated with inversion polymorphisms (reviewed in Smukowski & Noor, 2011) as different orientations of an inversion experience little successful recombination (reviewed in Kirkpatrick, 2010). Further research is warranted to characterize the nature of this differentiated region as well as whether it houses an inversion polymorphism.

While numerous “islands of differentiation” were observed between yellowhammers and pine buntings implying moderate genetic divergence, mtDNA introgression has the potential to homogenize the nuclear genomes of these taxa at mitonuclear genes by selecting for co-introgression of compatible alleles (Beck et al. 2015; Sloan et al. 2017; Morales et al. 2018). In the yellowhammer and pine bunting system, we found statistical support for the preferential introgression of mitonuclear genes into yellowhammers, but no statistical support for the preferential introgression into pine buntings. More specifically, our analysis showed that the proportion of introgressing mitonuclear genes was 1.7X higher in yellowhammers versus pine buntings. While these results suggest mitonuclear introgression primarily in the direction of pine buntings into yellowhammers, we also noted some consistency in the functions of putatively introgressing mitonuclear genes in both taxa.

Three of the mitonuclear genes within yellowhammer PIWs and three within pine bunting PIWs encode structural subunits of the ETC. The ETC is broken into five protein complexes which, through a series of enzymatic reactions, perform oxidative phosphorylation to produce ATP necessary for organism survival (reviewed in Ernster & Schatz, 1981). Four of the five ETC complexes are made up of subunits encoded by both the nuclear and mitochondrial genome (Hill, 2019) and correct fit between differentially encoded subunits is essential for the flow of electrons and protons across the ETC. To put this in perspective, changing even a single amino acid in one subunit can significantly disrupt its ability to interact with other subunits within an ETC complex (e.g. Gershoni et al. 2014). Because of the tight interactions within complexes and the consequences of subunit incompatibility, introgression of mtDNA is expected to select for co-introgression of mitonuclear genes encoding ETC structural subunits. Such co-introgression has been detected between differentially adapted populations of eastern yellow robin where mtDNA introgression between populations was followed by similar introgression of mitonuclear genes encoding subunits of complex I (Morales et al. 2018) and between different species of *Drosophila* where introgression and replacement of the mtDNA of one species during hybridization selected for co-introgression of genes that encode subunits of complex IV (Beck et al. 2015).

Of the ETC complexes, complex IV showed the strongest signal of co-introgression in the yellowhammer and pine bunting system. Three of the genes within yellowhammer PIWs and one gene within pine bunting PIWs were associated with this complex. As well, the gene COX5A—a structural subunit of complex IV—appeared in both sets of PIWs. It is unlikely that this gene introgressed in both directions, but it is possible that COX5A adaptively swept in both populations. In this situation, a particularly adaptive allele may have appeared in one species and swept to high frequency before co-introgressing into the other species following mtDNA introgression. Interestingly, the COX5A gene was one of the subunits that co-introgressed in the *Drosophila* example discussed above (Beck et al. 2015) lending some support to its particular importance to mitonuclear coevolution. More generally, complex IV is often used as a model for studying mitonuclear interactions due to its distinctive structure where a core of mitochondrial-encoded subunits is surrounded by nuclear-encoded subunits (Saraste, 1999). With such an excess of mitonuclear interactions, incompatibility involving complex IV has been investigated and detected in several systems including within primate xenomitochondrial cybrids (Barrientos et al. 2000) and between different species of *Drosophila* (Sackton et al. 2003). Furthermore, work by Osada & Akashi (2012) has provided strong evidence for compensatory coevolution between mitonuclear genes related to complex IV—including COX5A—and mtDNA among primates particularly at interacting amino acids of differentially encoded subunits. Altogether, these results suggest a crucial role for complex IV in mitonuclear coevolution as it may relate to divergence and speciation between yellowhammers and pine buntings.

Another group of mitonuclear genes that appeared consistently within the yellowhammer and pine bunting PIWs were those encoding subunits of the mitoribosome (five in yellowhammer PIWs and four in pine bunting PIWs). MRPS7, like COX5A, appeared in both yellowhammer and pine bunting PIWs suggesting that this gene may have adaptively swept through both taxa. Unlike the protein-protein interactions occurring within ETC complexes, mitonuclear interactions in the mitoribosome are between nuclear-encoded proteins and mitochondrial-encoded RNA (Hill, 2019). Protein subunits associate closely with rRNA during the formation of a mitoribosome, but also interact with mRNA and tRNA during the synthesis of mitochondrial proteins (Greber & Ban, 2016). Currently, research is limited on the extent and importance of interactions between mitoribosomal subunits and mitochondrial RNA. However, the fact that interactions between components are extensive and necessary for the synthesis of mitochondrial proteins suggests close coevolution between mtDNA and genes encoding mitoribosomal subunits that could strongly select for mitonuclear co-introgression following mtDNA introgression.

To summarize, yellowhammers and pine buntings are sister taxa that are divergent in appearance, song, and ecology (Panov et al. 2003; Rubtsov & Tarasov, 2017) yet vary greatly in their genomic differentiation from virtually none (at the mitochondrial genome) to nearly fixed (the differentiation peak on the Z chromosome). These patterns are best explained by a period of differentiation while geographically separated, followed by hybridization and mitochondrial DNA introgression. We observed evidence for preferential mitonuclear gene introgression (compared to introgression of other genes) from pine buntings into yellowhammers, as well as a tendency for mitonuclear genes encoding structural components of the ETC and the mitoribosome to appear within the PIWs of both taxa. One intriguing possibility is that mitonuclear co-introgression could have resulted in reduced mitonuclear incompatibilities between yellowhammers and pine buntings (Gershoni et al. 2009; Burton & Barreto, 2012; Hill, 2019), thereby contributing to their current extensive hybridization within central Siberia (Panov et al. 2003; 2007; Rubtsov, 2007; Rubtsov & Tarasov, 2017). This idea leads into the question— which can be addressed through a close examination of genomic variation within the hybrid zone—of whether the observed islands of differentiation on the Z and autosomes are sufficient in stabilizing yellowhammers and pine buntings as separate entities where they hybridize, or whether the two taxa are gradually merging into a single species.

## Author contributions

E. N., D.I., and A.S.R. conceived of this study. A.S.R. collected samples. E. N. and A.S.R. completed molecular techniques. E.G.M.N. conducted data analysis and wrote this manuscript with input from D.I. and A.S.R.

## Acknowledgements

For providing valuable feedback, we thank Dolph Schluter, Eric Taylor, Judith Mank, Elizabeth Natola, Rashika Ranasinghe, Kenneth Askelson, Finola Fogarty, Quinn McCallum, Ana Barreira, Jamie Clarke, Armando Geraldes and Jessica Irwin. For their kindness and support during field work, we thank the Tazeev family and Madelyn Ore. For providing additional samples, we thank The Bell Museum, The Burke Museum of Natural History and Culture, The Field Museum, The State Darwin Museum, The Swedish Museum of Natural History, The Zoological Museum of the Zoological Institute of the Russian Academy of Sciences, the Zoological Museum of the University of Copenhagen and their accompanying personnel. Major research funding was provided by the Natural Sciences and Engineering Research Council of Canada (NSERC CGSM award to E.N, Discovery Grants RGPIN-2017-03919 and RGPAS-2017-507830 awarded to D.I.) and by the Werner and Hildegard Hesse research awards (Research award in Ornithology and Fellowship in Ornithology awarded to E.G.M.N. by the University of British Columbia).

## Data Accessibility Statement

Raw DNA sequencing reads are available on the NBCI Sequence Read Archive (BioProject PRJNA768601). Read processing codes, barcodes, genotype data and R codes associated with statistical analyses will be made available on Dryad upon publication acceptance.

