## Supporting Information for "Extreme Heterogeneity in Genomic Differentiation between Phenotypically Divergent Songbirds: A Test of Mitonuclear Co-introgression"

**Supplementary Table 1.** Detailed information on all the samples included in this study. Where possible, museum accession numbers were included as part of the sample IDs following "Emberiza_GBS#_". Otherwise, samples were coded based on the museum they were obtained from or the collector's coding system. In the "sex" column, "m" stands for male, "f" stands for female and "uk" stands for unknown. The "TH" column contains the phenotypic scores of males for the throat plumage trait. The "BR" column contains the phenotypic scores of males for the brow plumage trait. The "BG" column contains the phenotypic scores of males for the background colour plumage trait. In the "Pheno. Class" (Phenotypic Class) column, the phenotypic classes of males are indicated as described in the methods section. Additional abbreviations include: "OUT" for outgroup, "FML" for female and "UK" for unknown. The numbers in the "Sampling Location" column correspond to those that appear in Figure 1A. In all columns, a "NA" observation stands for "Not Applicable".

| Sample ID | Species | Sex | TH | BR | BG | Pheno. Class | Source | Latitude (°N) | Longitude (°E) | Sampling Location |
|---|---|---|---|---|---|---|---|---|---|---|
| Emberiza_GBS1_ASR00_01 | E. leucocephalos | m | 7 | 7 | 7 | PL | Collected | 51.26 | 115.21 | 14 |
| Emberiza_GBS1_ASR05_05 | E. hortulana | uk | NA | NA | NA | OUT | Collected | NA | NA | NA |
| Emberiza_GBS1_ASR05_14 | E. citrinella | m | 1 | 2 | 1 | SC | Collected | 51.2 | 57.27 | 12 |
| Emberiza_GBS1_ASR05_17 | E. citrinella | m | 0 | 0 | 0 | PC | Collected | 51.2 | 57.27 | 12 |
| Emberiza_GBS1_ASR05_18 | E. citrinella | m | 2 | 1 | 1 | SC | Collected | 51.2 | 57.27 | 12 |
| Emberiza_GBS1_ASR05_33 | E. leucocephalos | m | 7 | 7 | 7 | PL | Collected | 51.12 | 118.56 | 15 |
| Emberiza_GBS1_ASR05_35 | E. leucocephalos | m | 7 | 7 | 7 | PL | Collected | 51.12 | 118.56 | 15 |
| Emberiza_GBS1_ASR05_36 | E. leucocephalos | m | 7 | 7 | 7 | PL | Collected | 51.12 | 118.56 | 15 |
| Emberiza_GBS1_ASR05_37 | E. leucocephalos | m | 7 | 7 | 7 | PL | Collected | 51.12 | 118.56 | 15 |
| Emberiza_GBS1_ASR05_43 | E. leucocephalos | m | 7 | 7 | 7 | PL | Collected | 51.12 | 118.56 | 15 |
| Emberiza_GBS1_ASR05_45 | E. leucocephalos | m | 7 | 7 | 7 | PL | Collected | 51.12 | 118.56 | 15 |
| Emberiza_GBS1_ASR05_47 | E. leucocephalos | m | 7 | 7 | 7 | PL | Collected | 51.12 | 118.56 | 15 |
| Emberiza_GBS1_ASR05_54 | E. leucocephalos | m | 7 | 7 | 7 | PL | Collected | 50.21 | 115.06 | 14 |
| Emberiza_GBS1_ASR05_55 | E. leucocephalos | m | 7 | 7 | 7 | PL | Collected | 50.21 | 115.06 | 14 |
| Emberiza_GBS1_ASR05_56 | E. leucocephalos | m | 7 | 7 | 7 | PL | Collected | 50.21 | 115.06 | 14 |
| Emberiza_GBS1_ASR05_59 | E. leucocephalos | m | 7 | 7 | 7 | PL | Collected | 50.21 | 115.06 | 14 |
| Emberiza_GBS1_ASR05_61 | E. leucocephalos | m | 7 | 7 | 7 | PL | Collected | 50.21 | 115.06 | 14 |
| Emberiza_GBS1_ASR05_66 | E. leucocephalos | m | 7 | 7 | 7 | PL | Collected | 50.21 | 115.06 | 14 |
| Emberiza_GBS1_ASR05_68 | E. leucocephalos | m | 7 | 7 | 7 | PL | Collected | 50.21 | 115.06 | 14 |
| Emberiza_GBS1_BKS_1572 | E. citrinella | m | 1 | 0 | 0 | SC | Burke Museum, USA | 54.573 | 39.182 | 7 |

| | | | | | | | | | | |
|---|---|---|---|---|---|---|---|---|---|---|
| Emberiza_GBS1_BKS_1646 | E. citrinella | f | NA | NA | NA | FML | Burke Museum, USA | 54.573 | 39.182 | 7 |
| Emberiza_GBS1_BKS_1821 | E. citrinella | m | 0 | 0 | 0 | PC | Burke Museum, USA | 51.412 | 34.547 | 6 |
| Emberiza_GBS1_BKS_1988 | E. citrinella | m | 0 | 0 | 0 | PC | Burke Museum, USA | 55.54 | 39.352 | 7 |
| Emberiza_GBS1_BKS_1989 | E. citrinella | m | 0 | 0 | 0 | PC | Burke Museum, USA | 55.54 | 39.352 | 7 |
| Emberiza_GBS1_BKS_2040 | E. citrinella | f | NA | NA | NA | FML | Burke Museum, USA | 55.54 | 39.352 | 7 |
| Emberiza_GBS1_BKS_2041 | E. citrinella | m | 1 | 0 | 0 | SC | Burke Museum, USA | 55.54 | 39.352 | 7 |
| Emberiza_GBS1_ENP97_25 | E. citrinella | uk | NA | NA | NA | UK | Collected | 58.33 | 44.76 | 11 |
| Emberiza_GBS1_EVN_327 | E. citrinella | uk | NA | NA | NA | UK | Bell Museum, USA | 61.45 | 38.67 | 8 |
| Emberiza_GBS1_EVN_331 | E. citrinella | m | 0 | 0 | 2 | SC | Bell Museum, USA | 61.45 | 38.67 | 8 |
| Emberiza_GBS1_EVN_357 | E. citrinella | uk | NA | NA | NA | UK | Bell Museum, USA | 61.45 | 38.67 | 8 |
| Emberiza_GBS1_EVN_363 | E. citrinella | uk | NA | NA | NA | UK | Bell Museum, USA | 61.45 | 38.67 | 8 |
| Emberiza_GBS1_EVN_573 | E. citrinella | f | NA | NA | NA | FML | Darwin Museum, Russia | 57.71 | 39.34 | 7 |

| | | | | | | | | | | |
|---|---|---|---|---|---|---|---|---|---|---|
| Emberiza_GBS1_EVN_574 | E. citrinella | m | 0 | 0 | 0 | PC | Darwin Museum, Russia | 57.71 | 39.34 | 7 |
| Emberiza_GBS1_IUK_615 | E. citrinella | m | 0 | 0 | 0 | PC | Bell Museum, USA | 61.45 | 38.67 | 8 |
| Emberiza_GBS1_IUK_631 | E. citrinella | f | NA | NA | NA | FML | Bell Museum, USA | 61.45 | 38.67 | 8 |
| Emberiza_GBS1_IVF_390 | E. citrinella | uk | NA | NA | NA | UK | Bell Museum, USA | 61.45 | 38.67 | 8 |
| Emberiza_GBS1_IVF_658 | E. cioides | uk | NA | NA | NA | OUT | Drovetsky expedition | NA | NA | NA |
| Emberiza_GBS1_IVF_682 | E. leucocephalos | m | 7 | 7 | 7 | PL | Drovetsky expedition | 50.50358 | 115.00286 | 14 |
| Emberiza_GBS1_M05_05 | E. citrinella | uk | NA | NA | NA | UK | Collected | 55.28 | 20.97 | 4 |
| Emberiza_GBS1_NVN_015 | E. citrinella | m | 0 | 0 | 0 | PC | Darwin Museum, Russia | 57.71 | 39.34 | 7 |
| Emberiza_GBS1_SVD_2134 | E. citrinella | m | 0 | 0 | 0 | PC | Burke Museum, USA | 43.54 | 40.47 | 9 |
| Emberiza_GBS1_SVD_2695 | E. citrinella | uk | NA | NA | NA | UK | Bell Museum, USA | 61.45 | 38.67 | 8 |
| Emberiza_GBS1_SVD_3479 | E. leucocephalos | m | 7 | 7 | 7 | PL | Drovetsky expedition | 49.64392 | 110.16517 | 13 |
| Emberiza_GBS1_SWM_03 | E. citrinella | uk | NA | NA | NA | UK | Swedish NHM | 57.99 | 12.49 | 1 |
| Emberiza_GBS1_SWM_10 | E. citrinella | uk | NA | NA | NA | UK | Swedish NHM | 65.86 | 21.48 | 5 |
| Emberiza_GBS1_SWM_12 | E. citrinella | uk | NA | NA | NA | UK | Swedish NHM | 65.86 | 21.48 | 5 |
| Emberiza_GBS1_VM_282a | E. leucocephalos | m | 7 | 7 | 7 | PL | Burke Museum, USA | 50.44 | 143.18 | 16 |

| | | | | | | | | | | |
|---|---|---|---|---|---|---|---|---|---|---|
| Emberiza_GBS2_ASR05_02 | E. citrinella | m | 1 | 0 | 0 | SC | Collected | 51.2 | 57.27 | 12 |
| Emberiza_GBS2_ASR05_15 | E. citrinella | m | 0 | 0 | 0 | PC | Collected | 51.2 | 57.27 | 12 |
| Emberiza_GBS2_ASR05_20 | E. citrinella | m | 2 | 1 | 0 | SC | Collected | 51.2 | 57.27 | 12 |
| Emberiza_GBS2_ASR05_22 | E. citrinella | m | 2 | 1 | 0 | SC | Collected | 51.2 | 57.27 | 12 |
| Emberiza_GBS2_ASR05_32 | E. leucocephalos | m | 7 | 7 | 7 | PL | Collected | 51.12 | 118.56 | 15 |
| Emberiza_GBS2_ASR05_38 | E. leucocephalos | m | 7 | 7 | 7 | PL | Collected | 51.12 | 118.56 | 15 |
| Emberiza_GBS2_ASR05_39 | E. leucocephalos | m | 7 | 7 | 7 | PL | Collected | 51.12 | 118.56 | 15 |
| Emberiza_GBS2_ASR05_40 | E. leucocephalos | m | 7 | 7 | 7 | PL | Collected | 51.12 | 118.56 | 15 |
| Emberiza_GBS2_ASR05_41 | E. leucocephalos | m | 7 | 7 | 7 | PL | Collected | 51.12 | 118.56 | 15 |
| Emberiza_GBS2_ASR05_44 | E. leucocephalos | m | 7 | 7 | 7 | PL | Collected | 51.12 | 118.56 | 15 |
| Emberiza_GBS2_ASR05_46 | E. leucocephalos | f | NA | NA | NA | FML | Collected | 51.12 | 118.56 | 15 |
| Emberiza_GBS2_ASR05_49 | E. leucocephalos | m | 7 | 7 | 7 | PL | Collected | 51.12 | 118.56 | 15 |
| Emberiza_GBS2_ASR05_51 | E. leucocephalos | m | 7 | 7 | 7 | PL | Collected | 50.21 | 115.06 | 14 |
| Emberiza_GBS2_ASR05_52 | E. leucocephalos | m | 7 | 7 | 7 | PL | Collected | 50.21 | 115.06 | 14 |
| Emberiza_GBS2_ASR05_57 | E. leucocephalos | m | 7 | 7 | 7 | PL | Collected | 50.21 | 115.06 | 14 |
| Emberiza_GBS2_ASR05_58 | E. leucocephalos | m | 7 | 7 | 7 | PL | Collected | 50.21 | 115.06 | 14 |
| Emberiza_GBS2_ASR05_60 | E. leucocephalos | m | 7 | 7 | 7 | PL | Collected | 50.21 | 115.06 | 14 |
| Emberiza_GBS2_ASR05_67 | E. leucocephalos | m | 7 | 7 | 7 | PL | Collected | 50.21 | 115.06 | 14 |
| Emberiza_GBS2_BKS_1583 | E. citrinella | m | 1 | 1 | 0 | SC | Burke Museum, USA | 54.573 | 39.182 | 7 |
| Emberiza_GBS2_BKS_1609 | E. citrinella | m | 1 | 0 | 0 | SC | Burke Museum, USA | 54.573 | 39.182 | 7 |
| Emberiza_GBS2_BKS_1654 | E. citrinella | m | 2 | 1 | 0 | SC | Burke Museum, USA | 51.349 | 37.123 | 6 |
| Emberiza_GBS2_BKS_1667 | E. citrinella | f | NA | NA | NA | FML | Burke Museum, USA | 51.349 | 37.123 | 6 |
| Emberiza_GBS2_BKS_2017 | E. citrinella | m | 1 | 0 | 1 | SC | Burke Museum, USA | 55.54 | 39.352 | 7 |

| | | | | | | | | | | |
|---|---|---|---|---|---|---|---|---|---|---|
| Emberiza_GBS2_EVN_328 | E. citrinella | uk | NA | NA | NA | UK | Bell Museum, USA | 61.45 | 38.67 | 8 |
| Emberiza_GBS2_EVN_366 | E. citrinella | uk | NA | NA | NA | UK | Bell Museum, USA | 61.45 | 38.67 | 8 |
| Emberiza_GBS2_EVN_570 | E. citrinella | m | 0 | 0 | 0 | PC | Darwin Museum, Russia | 57.71 | 39.34 | 7 |
| Emberiza_GBS2_F_12483 | E. cirlus | uk | NA | NA | NA | OUT | Collected | NA | NA | NA |
| Emberiza_GBS2_FMNH_01 | E. stewarti | uk | NA | NA | NA | OUT | Field Museum, USA AC# 347947 | NA | NA | NA |
| Emberiza_GBS2_IUK_2306 | E. aureola | uk | NA | NA | NA | OUT | Drovetsky expedition | NA | NA | NA |
| Emberiza_GBS2_IUK_2343 | E. leucocephalos | m | 7 | 7 | 7 | PL | Drovetsky expedition | 49.64392 | 110.16517 | 13 |
| Emberiza_GBS2_IUK_702 | E. citrinella | m | 0 | 0 | 0 | PC | Bell Museum, USA | 61.45 | 38.67 | 8 |
| Emberiza_GBS2_IUK_703 | E. citrinella | uk | NA | NA | NA | UK | Bell Museum, USA | 61.45 | 38.67 | 8 |
| Emberiza_GBS2_IUK_801 | E. citrinella | f | NA | NA | NA | FML | Bell Museum, USA | 65.85 | 44.24 | 10 |
| Emberiza_GBS2_M05_01 | E. citrinella | uk | NA | NA | NA | UK | Collected | 55.28 | 20.97 | 4 |
| Emberiza_GBS2_M05_03 | E. citrinella | uk | NA | NA | NA | UK | Collected | 55.28 | 20.97 | 4 |
| Emberiza_GBS2_M05_08 | E. citrinella | uk | NA | NA | NA | UK | Collected | 55.28 | 20.97 | 4 |
| Emberiza_GBS2_M05_10 | E. citrinella | uk | NA | NA | NA | UK | Collected | 55.28 | 20.97 | 4 |
| Emberiza_GBS2_SVD_1978 | E. calandra | uk | NA | NA | NA | OUT | Burke Museum, USA | NA | NA | NA |
| Emberiza_GBS2_SVD_3563 | E. leucocephalos | m | 7 | 7 | 7 | PL | Drovetsky expedition | 50.50358 | 115.00286 | 14 |

| | | | | | | | | | | |
|---|---|---|---|---|---|---|---|---|---|---|
| Emberiza_GBS2_SWM_11 | E. citrinella | uk | NA | NA | NA | UK | Swedish NHM | 59.81 | 17.05 | 2 |
| Emberiza_GBS2_VM_285a | E. leucocephalos | m | 7 | 7 | 7 | PL | Burke Museum, USA | 50.44 | 143.18 | 16 |
| Emberiza_GBS2_ZMUC_09 | E. citrinella | uk | NA | NA | NA | UK | ZMUC, Denmark | 51.71 | 18.61 | 3 |
| Emberiza_GBS4_F_13069 | E. cirlus | uk | NA | NA | NA | OUT | Collected | NA | NA | NA |
| Emberiza_GBS4_FM_347946 | E. stewarti | uk | NA | NA | NA | OUT | Field Museum, USA | NA | NA | NA |
| Emberiza_GBS4_K_Z2756 | E. cirlus | uk | NA | NA | NA | OUT | Collected | NA | NA | NA |
| Emberiza_GBS4_KZ_2766 | E. cirlus | uk | NA | NA | NA | OUT | Collected | NA | NA | NA |
| Emberiza_GBS4_MIM_165 | E. leucocephalos | f | NA | NA | NA | FML | Zoological museum, Russia | 50.68 | 142.97 | 16 |
| Emberiza_GBS4_RYA_2680 | E. leucocephalos | f | NA | NA | NA | FML | Zoological museum, Russia | 50.68 | 142.97 | 16 |
| Emberiza_GBS4_RYA_3178 | E. leucocephalos | m | NA | NA | NA | UK | Zoological museum, Russia | 50.68 | 142.97 | 16 |
| Emberiza_GBS4_SVN_2335 | E. leucocephalos | m | NA | NA | NA | UK | Zoological museum, Russia | 50.68 | 142.97 | 16 |
| Emberiza_GBS4_XD_657 | E. citrinella | m | 1 | 0 | 0 | SC | Collected | 56.06 | 36.13 | 7 |
| Emberiza_GBS4_XD_658 | E. citrinella | m | 0 | 0 | 1 | SC | Collected | 56.06 | 36.13 | 7 |
| Emberiza_GBS4_XD_659 | E. citrinella | m | 1 | 1 | 0 | SC | Collected | 56.06 | 36.13 | 7 |
| Emberiza_GBS5_ASR05_62_2 | E. leucocephalos | m | 7 | 7 | 7 | PL | Collected | 50.21 | 115.06 | 14 |
| Emberiza_GBS5_EAK_344 | E. stewarti | uk | NA | NA | NA | OUT | Zoological museum, Russia | NA | NA | NA |
| Emberiza_GBS5_F_13088 | E. cirlus | uk | NA | NA | NA | OUT | Collected | NA | NA | NA |
| Emberiza_GBS5_FM_347945 | E. stewarti | uk | NA | NA | NA | OUT | Field Museum, USA | NA | NA | NA |

| | | | | | | | | | | |
|---|---|---|---|---|---|---|---|---|---|---|
| Emberiza_GBS5_KZ_2762 | E. cirlus | uk | NA | NA | NA | OUT | Collected | NA | NA | NA |
| Emberiza_GBS5_RYA_3003 | E. leucocephalos | m | NA | NA | NA | UK | Zoological museum, Russia | 50.68 | 142.97 | 16 |
| Emberiza_GBS5_SVN_2336 | E. leucocephalos | f | NA | NA | NA | FML | Zoological museum, Russia | 50.68 | 142.97 | 16 |
| Emberiza_GBS5_XD_660 | E. citrinella | m | 1 | 1 | 0 | SC | Collected | 56.06 | 36.13 | 7 |
| Emberiza_GBS5_XD_661 | E. citrinella | m | 0 | 0 | 0 | PC | Collected | 56.06 | 36.13 | 7 |

**Supplementary Table 2.** Detailed information on the genomic locations and functions of the 162 mitonuclear genes investigated in this study. In the "Function" column, "ETC" stand for electron transport chain.

| Mitonuclear Gene | Chromosome | Start Position (bp) | End Position (bp) | Centre Position (bp) | Function |
|---|---|---|---|---|---|
| NDUFV3 | 1 | 4107093 | 4115372 | 4111232.5 | Structural subunit of ETC complex I |
| CARS2 | 1 | 24688325 | 24744232 | 24716278.5 | aminoacyl-tRNA synthetase |
| MRPS9 | 1 | 28543936 | 28583163 | 28563549.5 | Mitochondrial small ribosomal subunit protein |
| MRPL30 | 1 | 31234806 | 31237071 | 31235938.5 | Mitochondrial large ribosomal subunit protein |
| MRPL57 | 1 | 46619825 | 46621639 | 46620732 | Mitochondrial large ribosomal subunit protein |
| MRPS31 | 1 | 54773601 | 54793432 | 54783516.5 | Mitochondrial small ribosomal subunit protein |
| NARS2 | 1 | 86711128 | 86758715 | 86734921.5 | aminoacyl-tRNA synthetase |
| MRPL51 | 1 | 90281406 | 90282092 | 90281749 | Mitochondrial large ribosomal subunit protein |
| NDUFB4 | 1 | 92119879 | 92122800 | 92121339.5 | Structural subunit of ETC complex I |
| COX17 | 1 | 92525354 | 92527833 | 92526593.5 | Assembly factor/ancillary protein for ETC complex IV |
| MRPL48 | 1 | 98673934 | 98678389 | 98676161.5 | Mitochondrial large ribosomal subunit protein |
| TIMMDC1 | 1 | 103173445 | 103180367 | 103176906 | Assembly factor/ancillary protein for ETC complex I |
| MRPL39 | 1 | 112594984 | 112609353 | 112602168.5 | Mitochondrial large ribosomal subunit protein |
| ATP5J | 1 | 112654573 | 112658971 | 112656772 | Structural subunit of ETC complex V |
| GARS | 2 | 4436681 | 4465691 | 4451186 | aminoacyl-tRNA synthetase |
| MRPL32 | 2 | 32999048 | 33001488 | 33000268 | Mitochondrial large ribosomal subunit protein |
| FARS2 | 2 | 44068036 | 44292764 | 44180400 | aminoacyl-tRNA synthetase |
| MRPL3 | 2 | 62358147 | 62388654 | 62373400.5 | Mitochondrial large ribosomal subunit protein |
| LARS2 | 2 | 64965029 | 65050145 | 65007587 | aminoacyl-tRNA synthetase |
| MRPL36 | 2 | 91135578 | 91135946 | 91135762 | Mitochondrial large ribosomal subunit protein |
| NDUFS6 | 2 | 91137018 | 91144551 | 91140784.5 | Structural subunit of ETC complex I |
| NDUFV2 | 2 | 104093616 | 104113139 | 104103377.5 | Structural subunit of ETC complex I |
| MRPL15 | 2 | 116382079 | 116395598 | 116388838.5 | Mitochondrial large ribosomal subunit protein |
| TMEM70 | 2 | 124369603 | 124374001 | 124371802 | Assembly factor/ancillary protein for ETC complex V |
| MRPL53 | 2 | 127587659 | 127590834 | 127589246.5 | Mitochondrial large ribosomal subunit protein |

| Mitonuclear Gene | Chromosome | Start Position (bp) | End Position (bp) | Centre Position (bp) | Function |
|---|---|---|---|---|---|
| NDUFAF6 | 2 | 132905588 | 132922459 | 132914023.5 | Assembly factor/ancillary protein for ETC complex I |
| UQCRB | 2 | 133375437 | 133380089 | 133377763 | Structural subunit of ETC complex III |
| COX6C | 2 | 134936993 | 134941743 | 134939368 | Structural subunit of ETC complex IV |
| MRPL13 | 2 | 143508441 | 143536509 | 143522475 | Mitochondrial large ribosomal subunit protein |
| NDUFB9 | 2 | 145176891 | 145187833 | 145182362 | Structural subunit of ETC complex I |
| MRPS5 | 3 | 686280 | 701846 | 694063 | Mitochondrial small ribosomal subunit protein |
| NDUFAF5 | 3 | 5098403 | 5105436 | 5101919.5 | Assembly factor/ancillary protein for ETC complex I |
| MRPL33 | 3 | 7321655 | 7343178 | 7332416.5 | Mitochondrial large ribosomal subunit protein |
| IARS2 | 3 | 9631196 | 9663406 | 9647301 | aminoacyl-tRNA synthetase |
| MRPS10 | 3 | 14060720 | 14064492 | 14062606 | Mitochondrial small ribosomal subunit protein |
| MRPL2 | 3 | 19867530 | 19869125 | 19868327.5 | Mitochondrial large ribosomal subunit protein |
| PET117 | 3 | 23719042 | 23720746 | 23719894 | Assembly factor/ancillary protein for ETC complex IV |
| AARS2 | 3 | 31074744 | 31093395 | 31084069.5 | aminoacyl-tRNA synthetase |
| MRPL14 | 3 | 31303292 | 31314337 | 31308814.5 | Mitochondrial large ribosomal subunit protein |
| MRPS18A | 3 | 31771219 | 31792924 | 31782071.5 | Mitochondrial small ribosomal subunit protein |
| TFB2M | 3 | 33650775 | 33661865 | 33656320 | Mitochondrial Transcription Factor B2 |
| NDUFAF7 | 3 | 34173535 | 34180273 | 34176904 | Assembly factor/ancillary protein for ETC complex I |
| TFB1M | 3 | 54775521 | 54800918 | 54788219.5 | Mitochondrial Transcription Factor B1 |
| MRPL18 | 3 | 57591256 | 57593649 | 57592452.5 | Mitochondrial large ribosomal subunit protein |
| NDUFAF4 | 3 | 74973096 | 74976828 | 74974962 | Assembly factor/ancillary protein for ETC complex I |
| RARS2 | 3 | 78729006 | 78757996 | 78743501 | aminoacyl-tRNA synthetase |
| COX7A2 | 3 | 83154874 | 83158710 | 83156792 | Structural subunit of ETC complex IV |
| MRPL19 | 3 | 107287972 | 107290732 | 107289352 | Mitochondrial large ribosomal subunit protein |
| COX18 | 4 | 1462451 | 1466733 | 1464592 | Assembly factor/ancillary protein for ETC complex IV |
| MRPL1 | 4 | 2274392 | 2279731 | 2277061.5 | Mitochondrial large ribosomal subunit protein |
| MRPS18C | 4 | 8023084 | 8026575 | 8024829.5 | Mitochondrial small ribosomal subunit protein |
| NDUFC1 | 4 | 9943216 | 9945937 | 9944576.5 | Structural subunit of ETC complex I |

| Mitonuclear Gene | Chromosome | Start Position (bp) | End Position (bp) | Centre Position (bp) | Function |
|---|---|---|---|---|---|
| MRPL35 | 4 | 65701473 | 65703901 | 65702687 | Mitochondrial large ribosomal subunit protein |
| NDUFS8 | 5 | 7953211 | 7954951 | 7954081 | Structural subunit of ETC complex I |
| NDUFV1 | 5 | 8059469 | 8061462 | 8060465.5 | Structural subunit of ETC complex I |
| MRPL23 | 5 | 13894159 | 13905268 | 13899713.5 | Mitochondrial large ribosomal subunit protein |
| NDUFS3 | 5 | 21325493 | 21329936 | 21327714.5 | Structural subunit of ETC complex I |
| NDUFAF1 | 5 | 23445722 | 23451657 | 23448689.5 | Assembly factor/ancillary protein for ETC complex I |
| COX16 | 5 | 26716109 | 26760850 | 26738479.5 | Assembly factor/ancillary protein for ETC complex IV |
| NUBPL | 5 | 34372043 | 34456539 | 34414291 | Assembly factor/ancillary protein for ETC complex I |
| NDUFB1 | 5 | 45442861 | 45445862 | 45444361.5 | Structural subunit of ETC complex I |
| APOPT1 | 5 | 51884411 | 51890971 | 51887691 | Assembly factor/ancillary protein for ETC complex IV |
| TFAM | 6 | 4545977 | 4553892 | 4549934.5 | Mitochondrial Transcription Factor A |
| MRPS37/CHCHD1 | 6 | 14829186 | 14832429 | 14830807.5 | Mitochondrial small ribosomal subunit protein |
| NDUFB8 | 6 | 16430817 | 16434575 | 16432696 | Structural subunit of ETC complex I |
| COX15 | 6 | 21184712 | 21188712 | 21186712 | Assembly factor/ancillary protein for ETC complex IV |
| TWNK | 6 | 22511790 | 22514648 | 22513219 | mtDNA Helicase |
| NDUFA10 | 7 | 1914517 | 1940529 | 1927523 | Structural subunit of ETC complex I |
| ATP5G3 | 7 | 16650485 | 16653714 | 16652099.5 | Structural subunit of ETC complex V |
| NDUFS1 | 7 | 20857200 | 20871606 | 20864403 | Structural subunit of ETC complex I |
| NDUFB3 | 7 | 22144540 | 22146995 | 22145767.5 | Structural subunit of ETC complex I |
| MRPS14 | 8 | 105358 | 108090 | 106724 | Mitochondrial small ribosomal subunit protein |
| DARS2 | 8 | 2714489 | 2729279 | 2721884 | aminoacyl-tRNA synthetase |
| UQCRH | 8 | 18754346 | 18755566 | 18754956 | Structural subunit of ETC complex III |
| ATPAF1 | 8 | 19066784 | 19075132 | 19070958 | Assembly factor/ancillary protein for ETC complex V |
| MRPL37 | 8 | 22833154 | 22837987 | 22835570.5 | Mitochondrial large ribosomal subunit protein |
| PARS2 | 8 | 22937524 | 22940092 | 22938808 | aminoacyl-tRNA synthetase |
| MRPS22 | 9 | 680169 | 687341 | 683755 | Mitochondrial small ribosomal subunit protein |
| MRPL44 | 9 | 10644434 | 10649542 | 10646988 | Mitochondrial large ribosomal subunit protein |

| Mitonuclear Gene | Chromosome | Start Position (bp) | End Position (bp) | Centre Position (bp) | Function |
|---|---|---|---|---|---|
| NDUFB5 | 9 | 19985641 | 19991221 | 19988431 | Structural subunit of ETC complex I |
| COX5A | 10 | 1899415 | 1901700 | 1900557.5 | Structural subunit of ETC complex IV |
| POLG | 10 | 12982224 | 12992974 | 12987599 | Subunit of DNA polymerase gamma |
| MRPS11 | 10 | 13270849 | 13274501 | 13272675 | Mitochondrial small ribosomal subunit protein |
| MRPL46 | 10 | 13274531 | 13277811 | 13276171 | Mitochondrial large ribosomal subunit protein |
| COX4 | 11 | 21106 | 24901 | 23003.5 | Structural subunit of ETC complex IV |
| KARS | 11 | 12556178 | 12564535 | 12560356.5 | aminoacyl-tRNA synthetase |
| UQCRFS1 | 11 | 14269328 | 14270098 | 14269713 | Structural subunit of ETC complex III |
| ACAD9 | 12 | 1576425 | 1596683 | 1586554 | Assembly factor/ancillary protein for ETC complex I |
| UQCRC1 | 12 | 9243581 | 9252902 | 9248241.5 | Structural subunit of ETC complex III |
| NDUFAF3 | 12 | 12252246 | 12253315 | 12252780.5 | Assembly factor/ancillary protein for ETC complex I |
| MRPS25 | 12 | 21415500 | 21416915 | 21416207.5 | Mitochondrial small ribosomal subunit protein |
| UQCRQ | 13 | 1160234 | 1161216 | 1160725 | Structural subunit of ETC complex III |
| MRPL22 | 13 | 5404744 | 5411508 | 5408126 | Mitochondrial large ribosomal subunit protein |
| NDUFA2 | 13 | 15719950 | 15720141 | 15720045.5 | Structural subunit of ETC complex I |
| MRPS34 | 14 | 89067 | 91090 | 90078.5 | Mitochondrial small ribosomal subunit protein |
| NDUFB6 | 14 | 1409930 | 1411272 | 1410601 | Structural subunit of ETC complex I |
| MRPL28 | 14 | 2948403 | 2980093 | 2964248 | Mitochondrial large ribosomal subunit protein |
| EARS2 | 14 | 8805735 | 8811470 | 8808602.5 | aminoacyl-tRNA synthetase |
| NDUFAB1 | 14 | 8853879 | 8856325 | 8855102 | Structural subunit of ETC complex I |
| NDUFB10 | 14 | 9479824 | 9481800 | 9480812 | Structural subunit of ETC complex I |
| ATP5J2 | 14 | 11124929 | 11126597 | 11125763 | Structural subunit of ETC complex V |
| COX19 | 14 | 13640400 | 13642700 | 13641550 | Assembly factor/ancillary protein for ETC complex IV |
| UQCRC2 | 14 | 15520722 | 15531290 | 15526006 | Structural subunit of ETC complex III |
| MRPL40 | 15 | 8155635 | 8160277 | 8157956 | Mitochondrial large ribosomal subunit protein |
| COX6A1 | 15 | 10968121 | 10969778 | 10968949.5 | Structural subunit of ETC complex IV |
| UQCR10 | 15 | 12850007 | 12851340 | 12850673.5 | Structural subunit of ETC complex III |

| Mitonuclear Gene | Chromosome | Start Position (bp) | End Position (bp) | Centre Position (bp) | Function |
|---|---|---|---|---|---|
| MRPL41 | 17 | 1410903 | 1411829 | 1411366 | Mitochondrial large ribosomal subunit protein |
| SURF1 | 17 | 7671080 | 7676756 | 7673918 | Assembly factor/ancillary protein for ETC complex IV |
| MRPS2 | 17 | 8845099 | 8848425 | 8846762 | Mitochondrial small ribosomal subunit protein |
| NDUFA8 | 17 | 9921532 | 9923704 | 9922618 | Structural subunit of ETC complex I |
| MRPL12 | 18 | 1709716 | 1712644 | 1711180 | Mitochondrial large ribosomal subunit protein |
| POLG2 | 18 | 3399365 | 3405672 | 3402518.5 | Subunit of DNA polymerase gamma |
| COX10 | 18 | 3831022 | 3927582 | 3879302 | Assembly factor/ancillary protein for ETC complex IV |
| SCO1 | 18 | 5465970 | 5471599 | 5468784.5 | Assembly factor/ancillary protein for ETC complex IV |
| MRPL38 | 18 | 8209846 | 8218187 | 8214016.5 | Mitochondrial large ribosomal subunit protein |
| MRPS7 | 18 | 8759151 | 8762329 | 8760740 | Mitochondrial small ribosomal subunit protein |
| ATP5H | 18 | 8874306 | 8876875 | 8875590.5 | Structural subunit of ETC complex V |
| MRPL58 | 18 | 8888001 | 8890348 | 8889174.5 | Mitochondrial large ribosomal subunit protein |
| MRPL27 | 18 | 9156395 | 9159570 | 9157982.5 | Mitochondrial large ribosomal subunit protein |
| MRPS17 | 19 | 6083187 | 6086076 | 6084631.5 | Mitochondrial small ribosomal subunit protein |
| TTC19 | 19 | 7948708 | 7956750 | 7952729 | Assembly factor/ancillary protein for ETC complex III |
| ATP5E | 20 | 12072535 | 12073407 | 12072971 | Structural subunit of ETC complex V |
| MRPL20 | 21 | 4190134 | 4191696 | 4190915 | Mitochondrial large ribosomal subunit protein |
| MRPS38/AURKAIP1 | 21 | 4222177 | 4224571 | 4223374 | Mitochondrial small ribosomal subunit protein |
| NDUFS5 | 23 | 3948096 | 3948305 | 3948200.5 | Structural subunit of ETC complex I |
| MRPS15 | 23 | 5264268 | 5271412 | 5267840 | Mitochondrial small ribosomal subunit protein |
| ATP5L | 24 | 238825 | 240411 | 239618 | Structural subunit of ETC complex V |
| FOXRED1 | 24 | 7573063 | 7578176 | 7575619.5 | Assembly factor/ancillary protein for ETC complex I |
| TARS2 | 25 | 1192211 | 1197044 | 1194627.5 | aminoacyl-tRNA synthetase |
| ATP5F1 | 26 | 3213285 | 3216396 | 3214840.5 | Structural subunit of ETC complex V |
| MRPL45 | 27 | 4332672 | 4339508 | 4336090 | Mitochondrial large ribosomal subunit protein |
| ATP5G1 | 27 | 4567588 | 4568848 | 4568218 | Structural subunit of ETC complex V |
| NDUFA11 | 28 | 138984 | 139260 | 139122 | Structural subunit of ETC complex I |

| Mitonuclear Gene | Chromosome | Start Position (bp) | End Position (bp) | Centre Position (bp) | Function |
|---|---|---|---|---|---|
| UQCR11 | 28 | 1073915 | 1076925 | 1075420 | Structural subunit of ETC complex III |
| POLRMT | 28 | 2901392 | 2916002 | 2908697 | Mitochondrial RNA polymerase |
| NDUFS7 | 28 | 4121690 | 4125088 | 4123389 | Structural subunit of ETC complex I |
| ATP5D | 28 | 4237453 | 4239005 | 4238229 | Structural subunit of ETC complex V |
| ATP5C1 | 1A | 3456418 | 3463670 | 3460044 | Structural subunit of ETC complex V |
| NDUFA5 | 1A | 21570348 | 21574483 | 21572415.5 | Structural subunit of ETC complex I |
| MRPL42 | 1A | 44154725 | 44161681 | 44158203 | Mitochondrial large ribosomal subunit protein |
| NDUFA12 | 1A | 44613892 | 44626365 | 44620128.5 | Structural subunit of ETC complex I |
| NDUFA6 | 1A | 48813487 | 48815331 | 48814409 | Structural subunit of ETC complex I |
| YARS2 | 1A | 57351105 | 57356107 | 57353606 | aminoacyl-tRNA synthetase |
| SSBP1 | 1A | 58878378 | 58881966 | 58880172 | Single stranded DNA-binding protein |
| MRPS33 | 1A | 59199159 | 59217134 | 59208146.5 | Mitochondrial small ribosomal subunit protein |
| NDUFB2 | 1A | 59314052 | 59315170 | 59314611 | Structural subunit of ETC complex I |
| NDUFA9 | 1A | 63597529 | 63609489 | 63603509 | Structural subunit of ETC complex I |
| MRPS35 | 1A | 72765820 | 72780251 | 72773035.5 | Mitochondrial small ribosomal subunit protein |
| MRPS6 | 1B | 125717 | 175523 | 150620 | Mitochondrial small ribosomal subunit protein |
| ATP5O | 1B | 248176 | 252126 | 250151 | Structural subunit of ETC complex V |
| NDUFA1 | 4A | 9441670 | 9442588 | 9442129 | Structural subunit of ETC complex I |
| MARS2 | 4A | 16044198 | 16049852 | 16047025 | aminoacyl-tRNA synthetase |
| COX7B | 4A | 18172692 | 18174866 | 18173779 | Structural subunit of ETC complex IV |
| ATP5I | Z | 3439938 | 3442139 | 3441038.5 | Structural subunit of ETC complex V |
| MRPL50 | Z | 11148932 | 11151579 | 11150255.5 | Mitochondrial large ribosomal subunit protein |
| ATP5A1 | Z | 32785423 | 32792998 | 32789210.5 | Structural subunit of ETC complex V |
| MRPL17 | Z | 40019570 | 40020454 | 40020012 | Mitochondrial large ribosomal subunit protein |
| MRPS30 | Z | 44765927 | 44770194 | 44768060.5 | Mitochondrial small ribosomal subunit protein |
| NDUFS4 | Z | 46636290 | 46683735 | 46660012.5 | Structural subunit of ETC complex I |
| NDUFAF2 | Z | 49401620 | 49454084 | 49427852 | Assembly factor/ancillary protein for ETC complex I |

| Mitonuclear Gene | Chromosome | Start Position (bp) | End Position (bp) | Centre Position (bp) | Function |
|---|---|---|---|---|---|
| MRPS27 | Z | 65988920 | 66030873 | 66009896.5 | Mitochondrial small ribosomal subunit protein |
| COX7C | Z | 69702408 | 69704744 | 69703576 | Structural subunit of ETC complex IV |

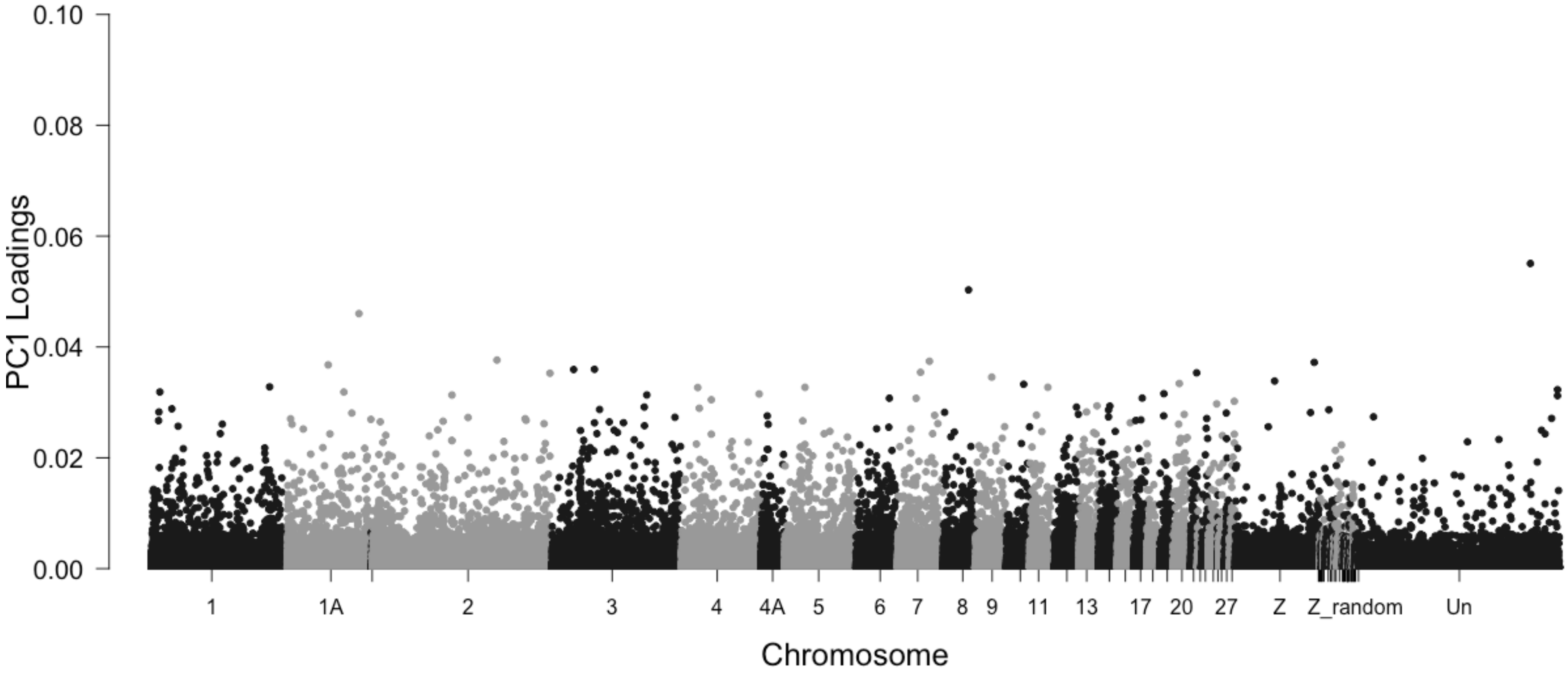


**Supplementary Figure 1.** PC1 loadings from the PCA shown in Figure 3 for 374,780 genome-wide SNPs plotted against genomic location if known. SNPs assigned to a "chromosome #_random" location are associated with a particular chromosome but their exact bp location is unknown. SNPs assigned to an "Un" location do not have any known location data.

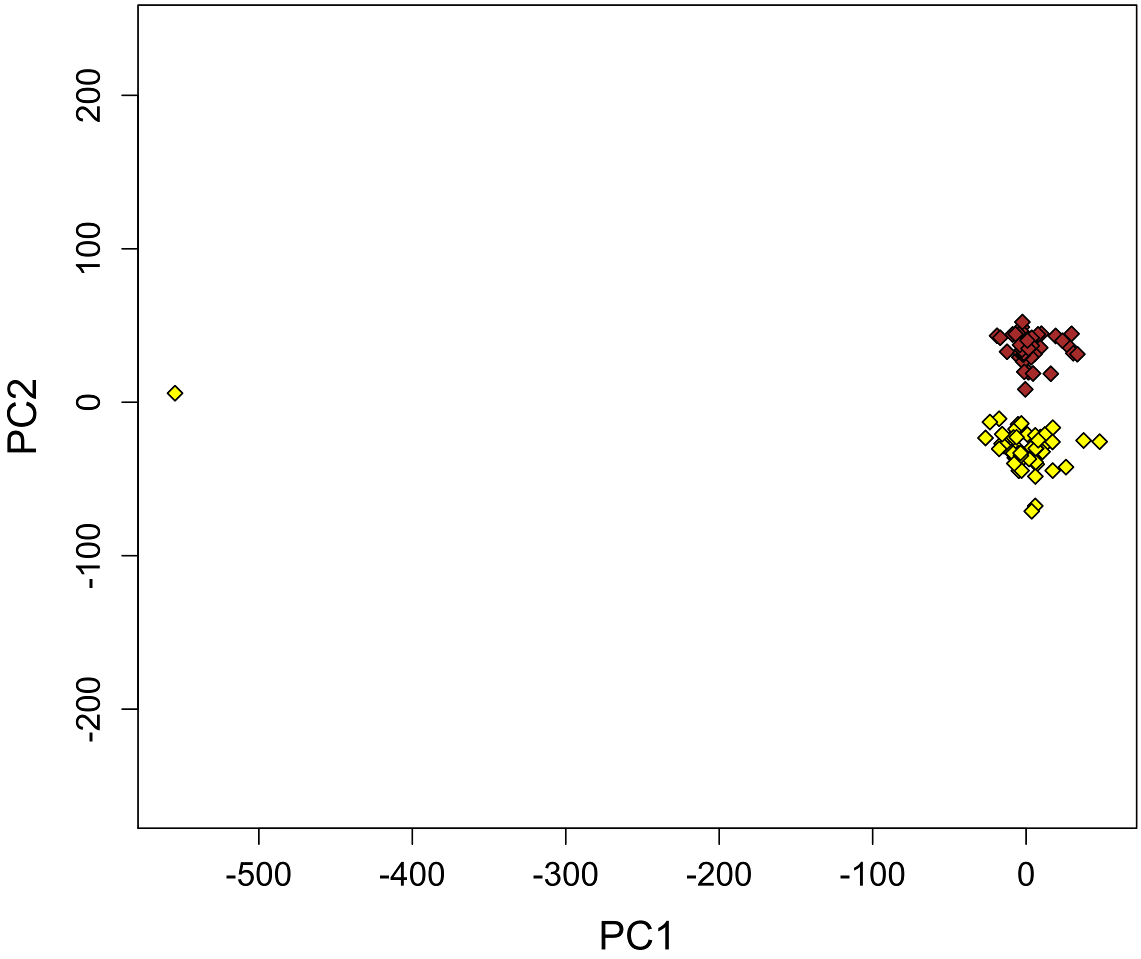


**Supplementary Figure 2.** PCA of genetic variation between allopatric yellowhammers (yellow; n = 53) and allopatric pine buntings (brown; n = 41) following the removal of sample "Emberiza_GBS2_ASR05_49". This sample was one of a pair of outliers that appeared in the PCA in Figure 3. PC1 explains 8.6% of the variation among individuals and PC2 explains 3.0% of the variation among individuals. Information from 374,780 genome-wide SNPs was included in this analysis.

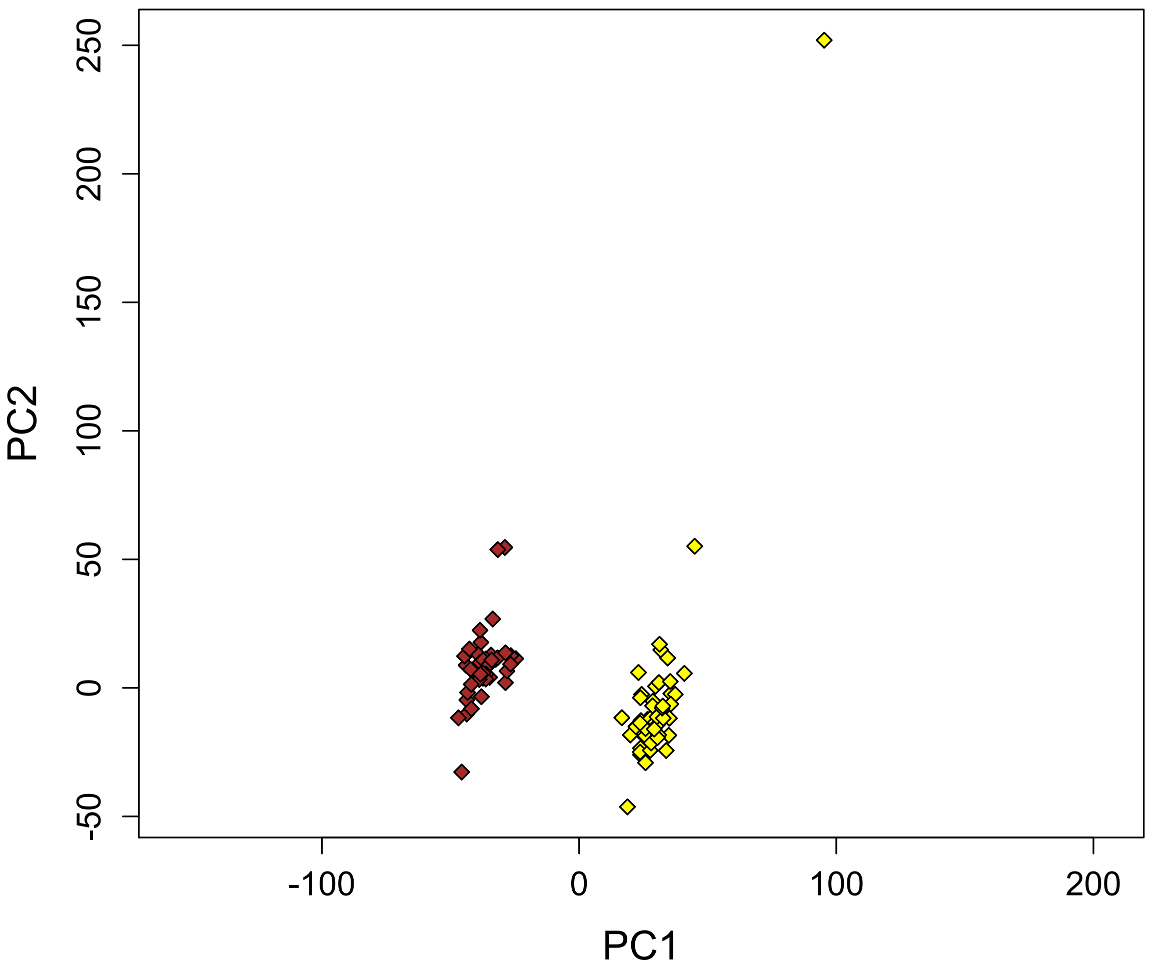


**Supplementary Figure 3.** PCA of genetic variation between allopatric yellowhammers (yellow; n = 52) and allopatric pine buntings (brown; n = 41) following the removal of samples "Emberiza_GBS2_ASR05_49" and "Emberiza_GBS2_BKS_1609". "Emberiza_GBS2_ASR05_49" was one of a pair of outliers that appeared in the PCA shown in Figure 3. "Emberiza_GBS2_BKS_1609" was an outlier that appeared in the PCA shown in Supplementary Figure 2. PC1 explains 3.2% of the variation among individuals and PC2 explains 2.6% of the variation among individuals. Information from 374,780 genome-wide SNPs was included in this analysis.

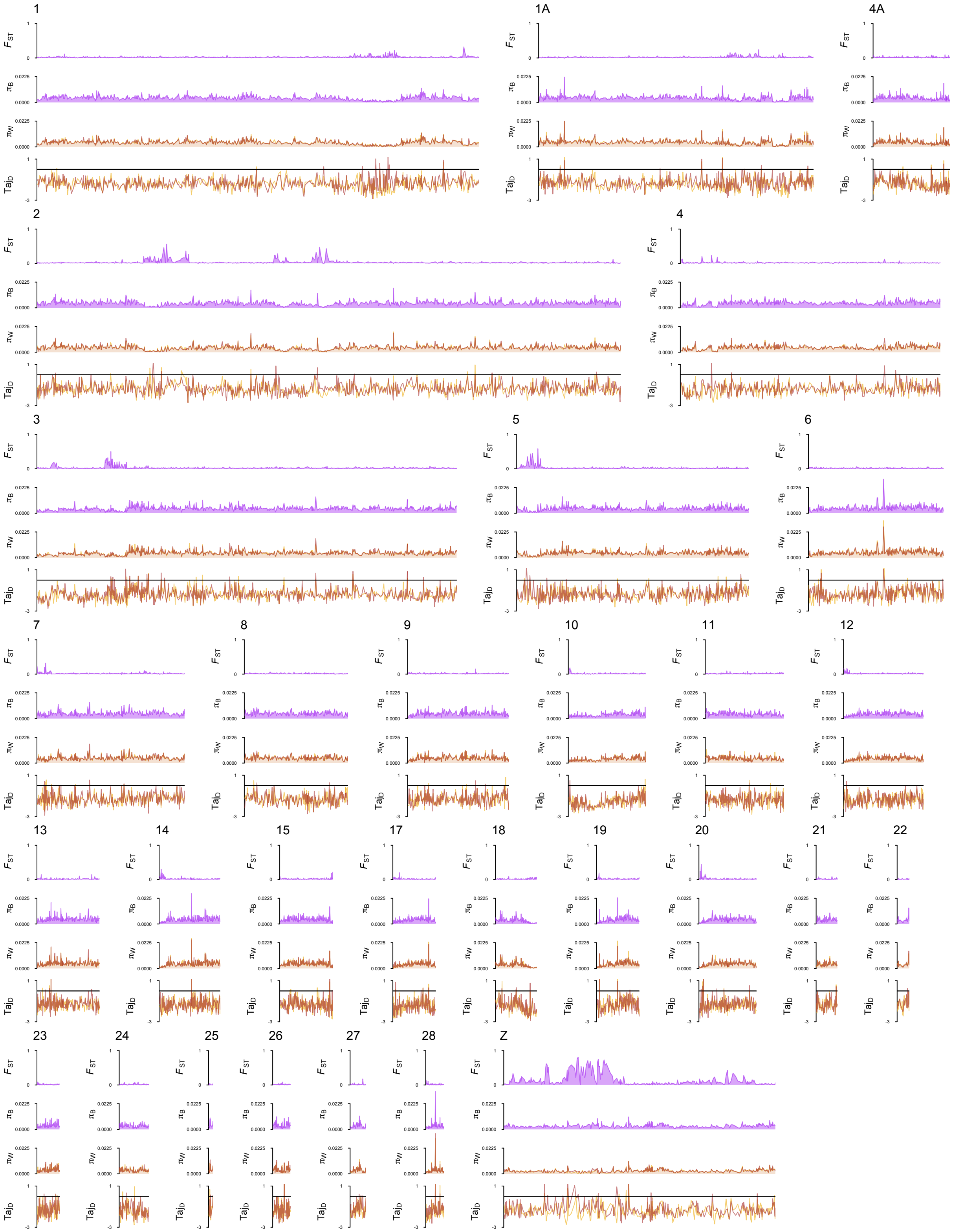


**Supplementary Figure 4.** Genome-wide patterns of genetic variation comparing allopatric yellowhammers (n = 53) and allopatric pine buntings (n = 42). Relative nucleotide differentiation

($F_{ST}$), absolute between-population nucleotide diversity ($\pi_B$), absolute within-population diversity ($\pi_W$) and Tajima's D ($Taj_D$) are shown as 2000 bp windowed averages across each chromosome. $F_{ST}$ and $\pi_B$ are shown as purple lines to indicate that values were calculated as a comparison between allopatric yellowhammers and pine buntings. $\pi_W$ and $Taj_D$ are shown as two separate lines (yellow = yellowhammers, brown = pine buntings) to indicate that values were calculated separately for each population
